# Individualized VDJ recombination predisposes the available Ig sequence space

**DOI:** 10.1101/2021.04.19.440409

**Authors:** Andrei Slabodkin, Maria Chernigovskaya, Ivana Mikocziova, Rahmad Akbar, Lonneke Scheffer, Milena Pavlović, Habib Bashour, Igor Snapkov, Brij Bhushan Mehta, Cédric R. Weber, Jose Gutierrez-Marcos, Ludvig M. Sollid, Ingrid Hobæk Haff, Geir Kjetil Sandve, Philippe A. Robert, Victor Greiff

**Author notes:** These authors jointly supervised this work (Philippe A. Robert, Victor Greiff).

## Abstract

The process of recombination between variable (V), diversity (D), and joining (J) immunoglobulin (Ig) gene segments determines an individual’s naïve Ig repertoire, and consequently (auto)antigen recognition. VDJ recombination follows probabilistic rules that can be modeled statistically. So far, it remains unknown whether VDJ recombination rules differ between individuals. If these rules differed, identical (auto)antigen-specific Ig sequences would be generated with individual-specific probabilities, signifying that the available Ig sequence space is individual-specific. We devised a sensitivity-tested distance measure that enables inter-individual comparison of VDJ recombination models. We discovered, accounting for several sources of noise as well as allelic variation in Ig sequencing data, that not only unrelated individuals but also human monozygotic twins and even inbred mice possess statistically distinguishable immunoglobulin recombination models. This suggests that, in addition to genetic, there is also non-genetic modulation of VDJ recombination. We demonstrate that population-wide individualized VDJ recombination can result in orders of magnitude of difference in the probability to generate (auto)antigen-specific Ig sequences. Our findings have implications for immune receptor-based individualized medicine approaches relevant to vaccination, infection, and autoimmunity.

## Introduction

The diversity, and thus recognition breadth, of adaptive immune receptor (AIR) repertoires (AIRR) is influenced by the recombination statistics of V, D, and J gene (and allele) segment recombination (Chi et al. 2020). Therefore, AIR recombination statistics determine the capacity to recognize antigens and to develop an adaptive immune response. Specifically, germline genes (and alleles), as well as their frequencies, have been linked to antibody neutralization breadth in infection (Avnir et al. 2016; Sangesland et al. 2020), the occurrence of precursor sequences of broadly neutralizing antibodies in the context of vaccine genetics (Lee et al. 2021) and autoantigen-specific binding in autoimmunity (Raposo et al. 2014; Parks et al. 2017).

With the advent of adaptive high-throughput AIRR sequencing (Weinstein et al. 2009), it has been observed that certain germline genes, and consequently recombined immune receptors, occur more often than others (Weinstein et al. 2009; Greiff et al. 2017a; Rubelt et al. 2016; Elhanati et al. 2018; Dupic et al. 2020). It has been shown that the occurrence of naïve AIRs could be predicted using a mathematical (explicit Bayesian or deep generative) model of VDJ recombination (Elhanati et al. 2018; Marcou et al. 2018; Olson and Matsen 2018; Davidsen et al. 2019) – hereafter called repertoire generation model (RGM). The Bayesian repertoire generation model parameters (RGMPs) correspond largely to those biological parameters that determine the biological mechanisms of VDJ recombination (Fig. 1A). Importantly, the RGMPs could be used to compute generation probabilities (Pgens) of a given AIR sequence. Although previous reports suggested that RGMP values differ across individuals (Briney et al. 2019; Marcou et al. 2018), the extent of this potential variation was neither quantified nor statistically verified. Inter-individual RGMP variation would imply that Pgens for identical AIR sequences differ across individuals. If this hypothesis is correct, it will implicate that each individual is restricted to exploring different AIR sequence spaces which again has implications for susceptibility to autoimmunity, cancer, and infectious diseases. For example, potentially important precursor AIR for vaccine responses (Lee et al. 2021; Sangesland et al. 2019) or potentially damaging autospecific AIR would occur more or less often depending on the individual’s RGMP.

**Figure 1:**
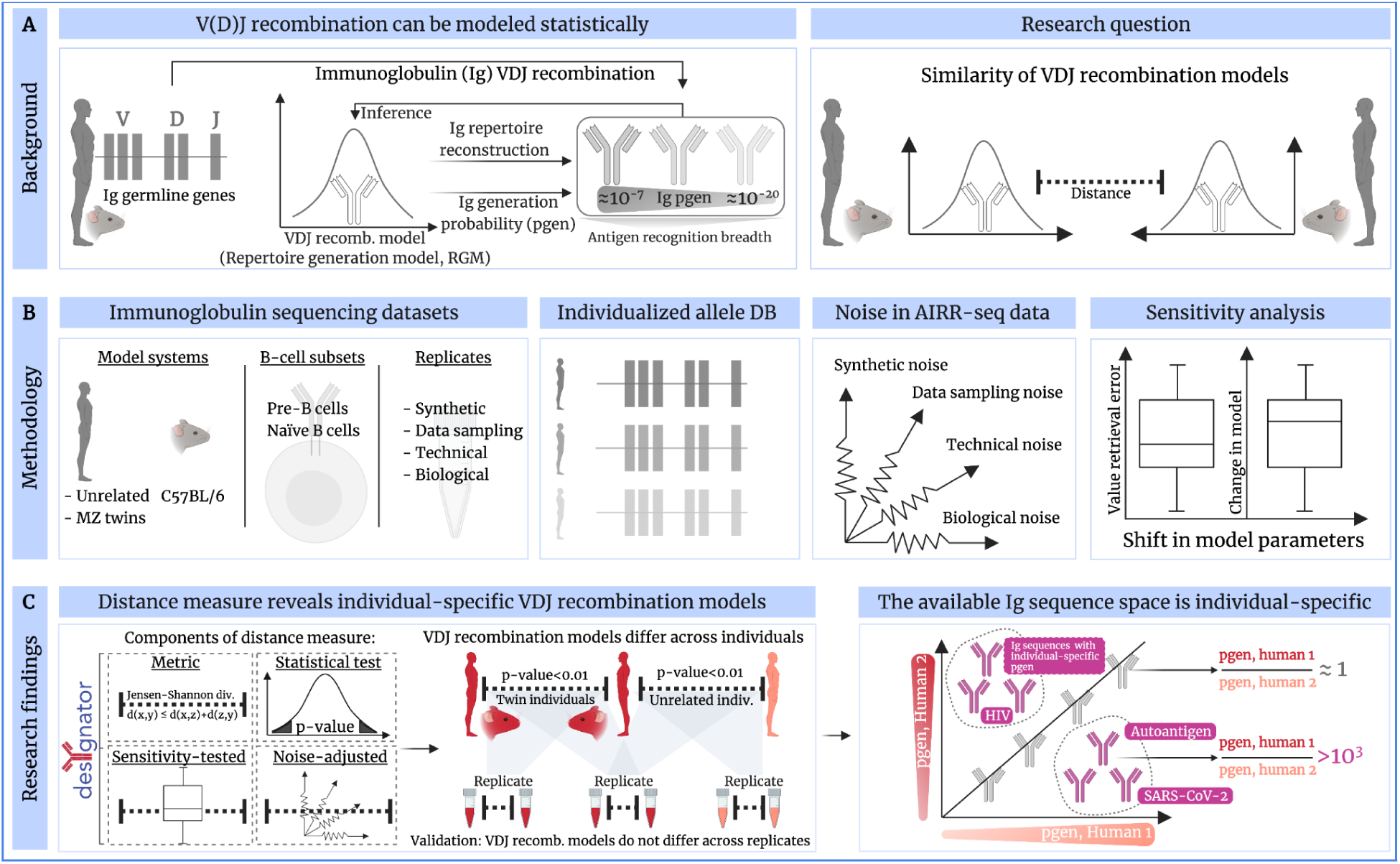
Comparison of immune receptor repertoire generation models. (A) The process of recombining variable (V), diversity (D), and joining (J) immunoglobulin (Ig) gene segments determines an individual’s naïve Ig repertoire, and consequently (auto)antigen recognition. VDJ recombination follows probabilistic rules that can be described statistically as repertoire generation models (RGMs). So far, it remains unknown whether VDJ recombination rules differ across individuals. We set out to resolve this question by developing a distance measure that enables the quantification of repertoire generation model parameter (RGMP) similarity. (B) Accounting for several sources of noise in murine and human Ig sequencing data (by leveraging various types of replicates) as well as allelic diversity, (C) we were able to implement a noise-aware, sensitivity-tested statistical test for comparing RGM similarity. We call our method desYgnator: DEtection of SYstematic differences in GeneratioN of Adaptive immune recepTOr Repertoires Using desYgnator, we found that replicate samples of the same subject are consistently more similar to each other than to samples from other unrelated individuals or even monozygotic twins (or inbred mice) indicating that not only genetic but also non-genetic factors contribute to the individualization of an RGM. We validated desYgnator by showing that RGM did not differ across synthetic and experimental replicates. We quantified the implication of individual RGMs on Ig repertoire architecture in a dataset of ≈100 human individuals by showing that the same (antigen-annotated) Ig sequence can have different generation probabilities across individuals. Thus, the available Ig sequence is individual-specific; predisposed by the individual RGM.

Here, we developed a novel computational and statistical approach (Fig. 1) for comparing RGMP sets (computed using IGoR; Marcou, Mora, and Walczak 2018). We call our method desYgnator (**DE**tection of **SY**stematic differences in **G**eneratio**N** of **A**daptive immune recep**TO**r **R**epertoires, Fig. 1C). We found that both unrelated and genetically identical test subjects (human monozygotic twins and inbred mice) had unique personal immunoglobulin RGMP values. Thus, an individual’s repertoire may not even be approximated by the RGM of this individual’s monozygotic twin. This suggests that non-genetic factors impact VDJ recombination. Our analysis of the VDJ recombination rules of twins versus non-twins allowed us to suggest a delineation of the specific regions of an Ig sequence that are subject to genetic heritability (Glanville et al. 2011). We validated this finding using synthetic and experimental replicates (Fig. 1C) and we accounted for common sources of noise in Ig-seq data as well as human germline allelic variation. We found that the effect of RGMP variation on the generation probability of Ig sequences, that were experimentally annotated with infectious disease and autoantigen specificity, may span several orders of magnitude. Thus, individualized VDJ recombination predisposes the available Ig sequence space (Fig. 1C). Our approach could be used to quantitatively study the effect of inter-individual differences in recombination statistics as a function of immune status, antigen recognition potential, and ethnicity (Avnir et al. 2016).

## Results

### A method for quantifying the similarity between repertoire generation models

Several studies have compared AIRRs across individuals using features such as germline gene usage (Bolen et al. 2017; Rubelt et al. 2016; Glanville et al. 2011), clonal overlap (Greiff et al. 2017a; Weinstein et al. 2009; Madi et al. 2014), clonal expansion (Stern et al. 2014; Greiff et al. 2015), and sequence similarity (Miho et al. 2018, 2019; Arora et al. 2018). However, all these features describe post VDJ recombination characteristics. So far, there is no sample-size-independent measure for comparing across individuals the entirety of RGMP, such as germline gene segment choice probabilities or deletion and insertion profiles (see Methods 5 for exact numbers of parameters).

Previously, Marcou and colleagues (Marcou et al. 2018) employed the Kullback-Leibler divergence (KLD, Kullback and Leibler 1951) for comparing the RGMP values inferred from a synthetic AIRR with ground truth RGMP values, i.e, those RGMP values used to generate that synthetic AIRR. The KLD was, however, not used to compare RGM from different individuals. In that case, the KLD decreased with increasing sample size, indicating, according to the authors, that the more sequencing reads were used for inference, the higher the inference precision. In this work, we favored using the Jensen-Shannon divergence (JSD), which is a smoothed symmetric version of the KLD. The square root of the JSD has the advantage of satisfying the triangle inequality (see Methods 5) enabling the computation of a relative distance between the RGMPs of any two repertoires (e.g., from different individuals or sub-samples of the same individual’s repertoire). The KLD is suited for quantifying by how much a distribution *P* diverges from a reference, perfectly known distribution *Q* (Marcou et al. 2018), while the JSD is designed to be used in a symmetric setting, i.e., with two arbitrary distributions. The JSD was previously also employed by Sethna and colleagues (Sethna et al. 2020) for comparing TCR RGMPs inferred from 651 individuals with a “universal” RGM (inferred from TCR sequences randomly drawn from all individual samples). However, this study did not compare individual RGMP sets in a pairwise fashion.

As various sources of noise can arise in AIRR data (Puelma Touzel et al. 2020; Koraichi et al. 2021), we aimed to quantify their impact on the pairwise JSD of RGMPs. Noise quantification depends on suitable replicates. Therefore, we examined the selected potential sources of sample-associated noise by means of different types of replicates (Fig. 2):

(i) synthetic replicates (cyan curves): synthetic samples generated using the same set of IGoR parameter values. These samples differ only by the unavoidable *synthetic sampling noise* and thus allow for its quantification.
(ii) data replicates (pale blue curves) subsampled without replacement from the same AIRR FASTA file; these replicates differ only by the *data sampling noise*.
(iii) technical replicates (solid blue curves) for which the RNA samples were split and then sequenced independently; these samples differ due to the *technical noise,* and data sampling noise. The MOUSE_PRE and MOUSE_NAIVE datasets each contain a pair of technical replicates (see Methods 1).
(iv) biological replicates (violet curves) obtained from the same individual and differ due to the *biological noise* (subsampling of the B/T cells from the actual repertoire, different expression levels), as well as due to both technical noise and data sampling noise. The HUMAN1 dataset contains a single pair of biological replicates.

**Figure 2:**
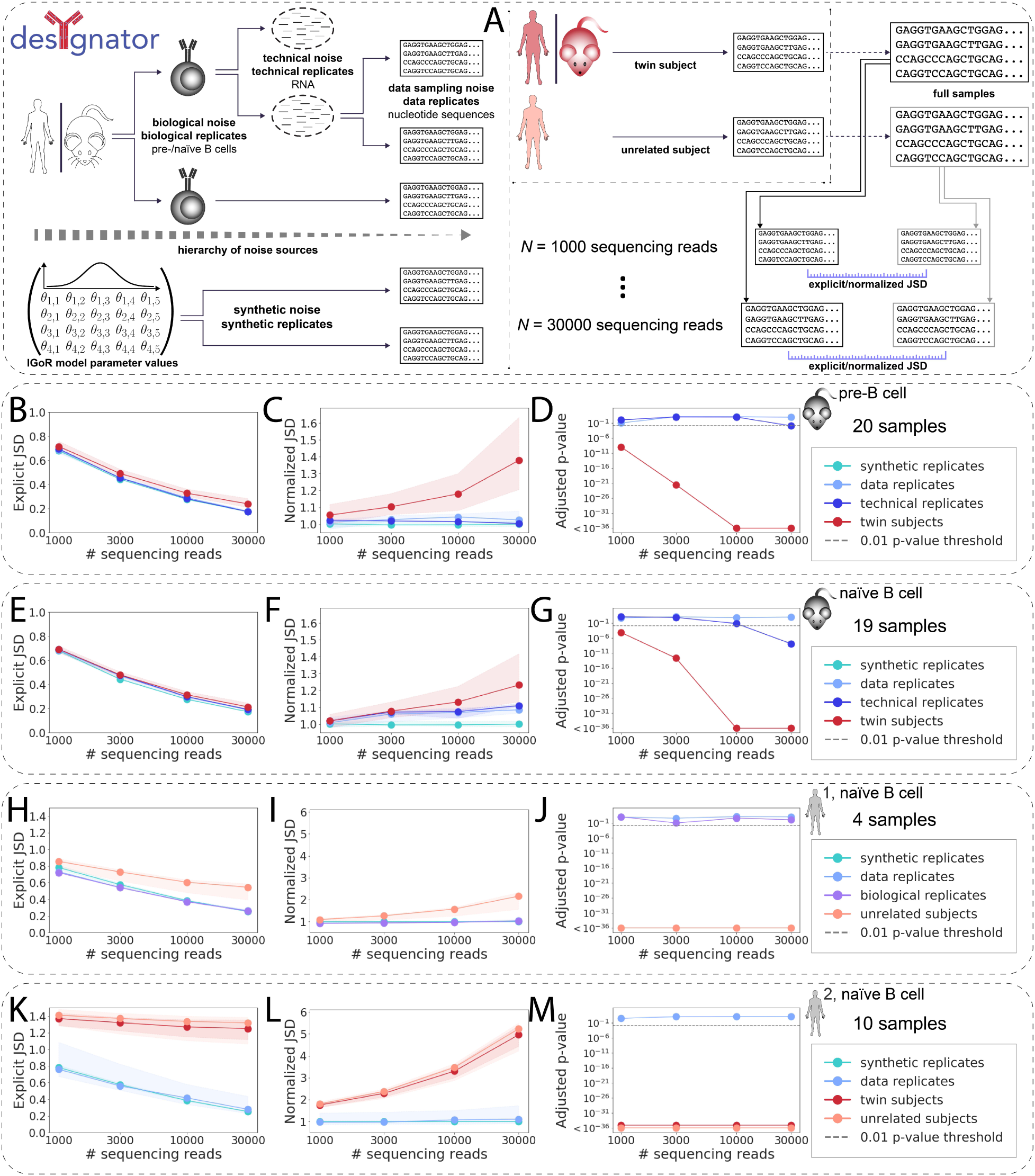
Repertoire generation model parameters are individual-specific independently of the degree of immunogenetic similarity between individuals. (A) Different sources of AIRR-seq noise may arise impacting repertoire generation model parameter (RGMP) inference. To account for these sources of noise, different kinds of replicates are necessary. Specifically, biological replicates (i.e., biological samples obtained from the same individual) allow for observing **biological noise**; technical replicates (an RNA sample that was split and the parts were sequenced independently) allow for observing **technical noise;** data replicates (subsamples of the same AIRR FASTA file) allow for observing **data sampling noise**. Samples obtained from different (either twin or unrelated) subjects incorporate all these aforementioned sources of noise along with the associated potential non-genetic or genetic individual differences between their RGMPs. Synthetic replicates (synthetic samples generated using the same RGMP) allow for observing **synthetic noise**. (B) Explicit JSD (Jensen-Shannon divergence) between RGMP inferred from samples differing by several levels of noise: *synthetic* replicates; *data* replicates; *technical* replicates; *twin* mice. We computed the explicit JSD for random subsets of [1000, 3000, 10000, 30000] sequencing reads taken from samples of the MOUSE_PRE dataset (nineteen IgH pre-B cell samples from C57BL/6 mice and one technical replicate, see Methods 1). Circles correspond to the median explicit JSD, shaded areas correspond to the whole range of the explicit JSD for the given sample size and pair type (from minimum to maximum). (C) The amount of noise that accounts for the difference between synthetic replicates is quantified using the explicit JSD. This can be considered as the lower bound of noise in our system. We then normalized the explicit JSD by this lower bound. (D) To test whether the difference between a pair of samples is significantly higher than the difference between data replicates, we adapted the Student’s T-test. The adjusted p-values for data and technical replicates were above the 0.01 threshold for each sample size except 30000 for technical replicate. The adjusted p-values for twin subjects were below the 0.01 threshold for all sample sizes, indicating that the recombination models of the twin subjects are not identical. (E–G) Same as B–D but computed for the MOUSE_NAIVE dataset (nineteen IgH naïve B cell samples from C57BL/6 mice and one technical replicate). The twin subjects are closer to each other than in the pre-B cell case. The p-values of the statistical test, as in D, indicated RGMP of cross-subject samples differed systematically. (H–J) Same as B–D but computed for the HUMAN1 dataset (three IgH naïve B cell samples of healthy Caucasian male donors and one biological replicate). For all samples, individually restricted germline allele databases were constructed. The considered sample pair types are: *synthetic* replicates; *data* replicates; *biological* replicates; *unrelated* subjects. P-values indicate that biological as well as technical replicates were generated with the same RGMPs and that RGMPs differed across unrelated human individuals. (K–M) Same as B–D but computed for the HUMAN2 dataset (IgH naïve B cell samples from five pairs of monozygotic twins). For all samples, individually restricted germline allele databases were constructed. The considered sample pair types are: *synthetic* replicates; *data* replicates; *twin* subjects; *unrelated* subjects. P-values indicate that RGMPs of human monozygotic twins differ. All p-values were adjusted using the Bonferroni correction within one dataset. The significance threshold of p=0.01 is indicated by a grey line.

Samples from human monozygotic twin subjects and inbred mice (red curves, Fig. 2), and samples from unrelated human subjects (tan curves, Fig. 2), both incorporate all three sources of noise (not including synthetic noise). In addition, they incorporate potential non-genetic (measurable in monozygotic twins and inbred mice) or genetic systematic differences.

Comparing the distance between twin subjects with the distance between unrelated subjects allows for the discrimination between non-genetic and genetic factors underlying the systematic differences in AIRR generation. Here, we developed a method for detecting such differences, **DE**tection of **SY**stematic differences in **G**eneratio**N** of **A**daptive immune recep**TO**r **R**epertoires (**desYgnator**).

To quantitatively discriminate noise from systematic differences, we first investigated how the JSD (hereafter: ‘explicit JSD’) is impacted by each level of noise with sample sizes from 1000 to 30000 sequencing reads (see Methods 4 for the definition of sequencing read and Methods 5 for an explanation of sample size) on murine (Fig. 2B and E, MOUSE_PRE and MOUSE_NAIVE datasets) and human (Fig. 2H and K, HUMAN1 and HUMAN2 datasets) samples.

Consistent with the KLD observations by Marcou and colleagues (Marcou et al. 2018), we found that the explicit JSD gradually decreases with an increasing number of sequencing reads (Fig. 2B,E,H,K). Explicit JSD values for data replicates, technical replicates, and biological replicates were similar suggesting that the noise introduced by the technological processes (and even by the biological processes, such as different expression levels) was negligible compared to the data sampling noise.

However, models inferred from samples of 30000 sequencing reads obtained from different subjects were all closer to each other (i.e., the explicit JSD between them was lower) than to models inferred from samples of 3000 sequencing reads obtained from the same subject (Fig. 2B,E,H,K). Thus, the explicit JSD between inferred RGMPs is sample size-dependent (i.e. the RGMP estimates are skewed differently for different sample sizes), suggesting that either the inferred parameters are different, or that the models have different levels of complexity. Therefore, it is not recommended to directly compare RGMP inferred from samples of different *sizes.* Furthermore, the threshold for determining the statistically significant pairwise difference of RGMP sets is also sample size-dependent and thus may vary across sample sizes.

To compensate for this dependence of the thresholds on the sample size, we introduced a normalized JSD (see Methods 5), which is obtained by dividing the explicit JSD of two RGMPs by the average explicit JSD between synthetic replicates (of the same sample size). For the normalization, we computed the explicit JSD between fifteen independently generated pairs of synthetic replicates (we used RGMP inferred from the first sample of the HUMAN2 dataset to generate human synthetic replicates and RGMP inferred from the first sample of the MOUSE_PRE dataset to generate mouse synthetic replicates). To validate that JSD normalization allows compensating for potential dependencies of the explicit JSD on sample size, we repeated all explicit JSD calculations for the normalized JSD (Fig. 2C,F,I,L).

The normalized JSD, unlike the explicit one, followed a clear pattern: in the cases where the underlying RGMP were assumed (and supposed) to be identical, i.e., for all types of replicates representing the same individual (Fig. 2C,F,I,L), the normalized JSD remained on the same level with increasing sample size (cyan, pale blue, solid blue, and violet). On the contrary, it increased for the samples obtained from unrelated (Fig. 2I,L) and even from twin subjects (Fig. 2C,F,L), tan and red respectively. This showed that the RGMP difference between individuals is distinguishable from the above-mentioned levels of noise provided the number of sequencing reads is sufficiently high, and that the normalized JSD, unlike the explicit JSD, enables the detection of this difference via a sample size-independent threshold. Of note: we expect the normalized JSD to stabilize for a high enough sample size.

To investigate the identifiability of the inter-individual RGMP difference, we derived a statistical test (see Methods 6) that compares the JSD between two samples (technical or biological) to the expected level of noise from data replicates, and showed the associated p-values in panels (Fig. 2D,G,J,M), related to panels (Fig. 2C,F,I,L) respectively. Altogether, using the developed statistical test (see Methods 6) we failed to reject the null hypothesis that the explicit and normalized JSD between RGMP sets of technical (for a sample size of [1000, 3000, 10000] sequencing reads) or biological (for all considered sample sizes) replicates was higher than the difference between RGMP sets of data replicates for all samples. This supports the statement that both technical and biological noise was in most cases dominated by the data sampling noise. When we tested the homozygous twins and unrelated subjects in the same way against the data replicates, there was a statistically significant difference (and even for the sample size of 30000 reads, the technical noise was negligible when compared to the inter-individual difference). The test showed that 1000 sequencing reads are sufficient to observe a significant difference (adjusted p-value < 0.01) in the case of murine pre-/naïve B cell data and human naïve B cell data (Fig. 2D,G,J,M).

To measure the impact of genetic factors on the normalized JSD, we computed the pairwise normalized JSD for five pairs of monozygotic twins (HUMAN2 dataset, Fig. S9). We found that the normalized JSD between the RGMPs of samples obtained from twin subjects is on average 5% lower than between samples obtained from unrelated subjects, which may indicate that non-genetic factors account for approximately 90% of the normalized JSD difference (taking into account that some of the normalized JSD is due to the noise). These data support the view that non-genetic factors play an important role in the AIRR generation, which was also noted for T cell repertoires (Dupic et al. 2020).

To explore the impact of different components of the RGM (e.g., V segment choice probability, V deletions, J given V choice conditional probabilities, see Methods 5) on the explicit and normalized JSD between the models, we reproduced the afore-described experiment using only V-choice-agnostic parameters (i.e., excluding IGHV-related parameters and only investigating: J choice, J deletion, D choice, D deletion, VD insertion, DJ insertion, Fig. S5, columns 2 and 4) and only V-related parameters (V choice, V deletion, Fig. S5, columns 1 and 3). In both cases, the data validate the results obtained with the full model except that for the V-choice-agnostic models, where the distances between samples from twin human subjects were not lower than the distances between samples from unrelated human subjects. This may indicate that genetic (genetically heritable) factors are only responsible for the difference in V-related RGMP – and, consequently, the difference in other RGMP (J segment choice, D segment choice, insertion and deletion profiles) is entirely caused by non-genetic factors. This is consistent with the previous findings for post-recombination statistics in AIRR-seq samples, i.e. that only V segment usage is genetically heritable and not D or J segment usage, or CDR3 similarity (Glanville et al. 2011; Rubelt et al. 2016).

The technological process and the data preprocessing workflow may introduce unavoidable bias into AIRR-seq sample generation (for example, different choices of PCR primers may result in very different sequenced repertoires). To explore the influence of this bias on RGMP inference, we compared the average values of the normalized JSD for the HUMAN1 (where the sequences were generated with Illumina HiSeq: 2×125bp – shorter reads, high sequencing depth) and HUMAN2 (Illumina MiSeq: 2×300bp – longer reads, lower sequencing depth) datasets (Fig. 2I and L). While we are able to compare distances between human models that have the same parameters (same model structure), it is not possible to compare murine and human datasets as their models represent different sets of V, D and J segments. The average normalized JSD differed two-fold between the HUMAN1 and HUMAN2 datasets for the sample size of 30000 sequencing reads, indicating the presence of a technological bias in RGMPs. Thus, caution and care are advised when comparing IGoR models inferred from samples generated using different experimental protocols.

To conclude, using the normalized JSD, we found that IgH RGMP differed not only across unrelated individuals but also across inbred C57BL/6 mice and human monozygotic twins, which may indicate that the rules of immune receptor recombination are governed not only by genetic factors but also by non-genetic ones.

### Immunoglobulin repertoire generation model parameters are unique across individuals

To explore the variation of VDJ recombination rules on a larger scale in human individuals (in naïve B-cell repertoires), we computed the normalized JSD on a cohort of 99 unrelated individuals from the HUMAN3 dataset (Gidoni et al. 2019 see Methods 1, Fig. 3A). Analogously to the HUMAN1 and the HUMAN2 datasets (Fig. 2), we constructed individually restricted germline allele databases for all samples via merging the RGMPs corresponding to alleles of the same V/D/J gene and calculated the explicit and normalized JSD on the gene-level RGMPs (see Methods 2). For the sample size of 10000 and 30000 sequencing reads, all pairwise distances were higher than the distance between data replicates, indicating that again all individuals had different RGMPs. For lower sample sizes of 1000 and 3000 sequencing reads, the distances were comparable to those between data replicates. This suggests that at least 10000 sequencing reads are required to overcome noise when building personalized RGMs. Analogous results were obtained when investigating the IGHV-only and IGHV-agnostic explicit/normalized JSD (Fig. S7 and Fig. S8).

**Figure 3:**
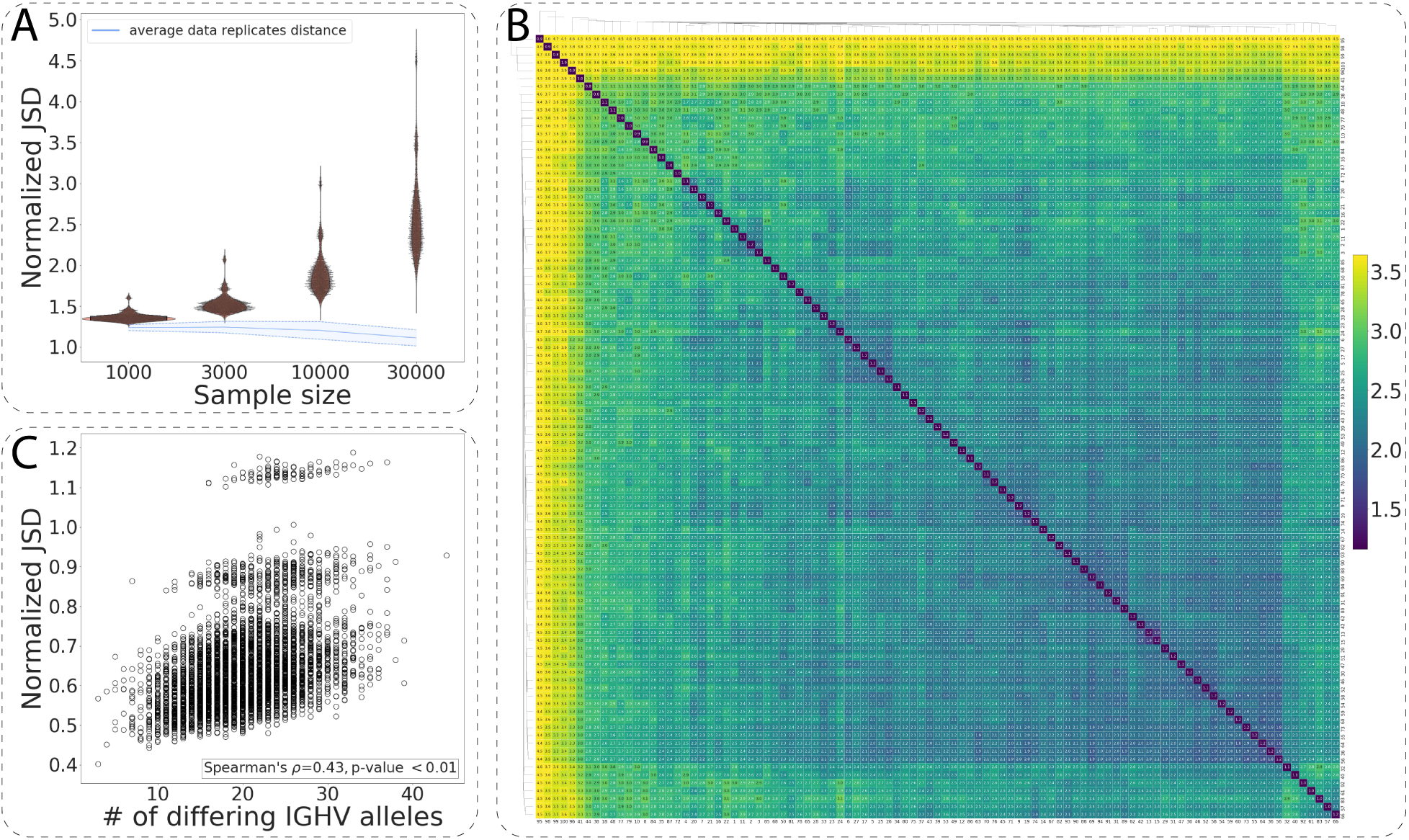
Immunoglobulin repertoire generation model parameters are unique across human individuals. (A) The distribution of the pairwise normalized JSD for 99 individuals of the HUMAN3 dataset computed for subsamples of 1000, 3000, 10000 and 30000 sequencing reads. The blue line corresponds to the average distance between data replicates. (B) Heatmap visualization of (A) for the subsample size of 30000 sequencing reads: the values on the diagonal correspond to the average distance between data replicates. (C) The number of IGHV gene alleles that differ between any two individuals as a function of the normalized JSD between their RGMP inferred from subsamples of 30000 sequencing reads.

To visualize the similarity variation of RGMP in the HUMAN3 dataset, we performed hierarchical clustering (single-linkage clustering, Müllner 2011) on pairwise computed normalized JSD (Fig. 3B). Although the range of the normalized JSD was large (min: 1.59, max: 4.71, median: 2.49), there were no clear clusters. Of note, we detected a moderate correlation of pairwise differences in RGMP values with IGH allele repertoire similarity (Fig. 3C, Spearman’s correlation coefficient = 0.43, p-value < 10^-36^, see also Fig. S7 and Fig. S8), suggesting that the difference in RGMP values between individuals may in part be explained by IGHV gene polymorphisms.

### Generation probabilities of antigen-annotated immunoglobulin sequences vary within and among related and unrelated human individuals

A direct corollary of the variation of RGMPs across individuals is the variation of individualized generation probabilities (Pgen) for the same Ig sequence. To study and quantify Pgen variation, we analyzed generation probabilities (as computed by the RGMs corresponding to individuals from the HUMAN2 and HUMAN3 datasets) of Ig sequences with known antigen specificity.

We assembled antigen-annotated Ig data from three sources (Swindells et al. 2017; Raybould et al. 2020; Roy et al. 2017) leading to 3492 unique CDRH3 (hereafter only referred to as CDR3) amino acid sequences specific to seven antigens in total: SARS-CoV2 (1062 sequences), Transglutaminase 2 (or TG2 [autoantigen], 1048 sequences), HIV (324 sequences), Tetanus (290 sequences), Influenza (283 sequences), MERS-CoV (97 sequences), SARS-CoV1 (84 sequences). We also included 304 sequences that were specific to both SARS-CoV1 and SARS-CoV2. To calculate the Pgens of CDR3 amino acid sequences, we used OLGA (Sethna et al. 2019) based on the RGMPs computed with IGoR for Fig. 2 and Fig. 3 (all RGMP were inferred from samples of 30000 sequencing reads).

To analyze the consistency of the Pgens across the RGMPs inferred from different samples, we first used the models corresponding to the individuals from the HUMAN2 dataset (Fig. 4A): a model inferred from the sample obtained from the pair 1 twin A individual, a model inferred from a data replicate sample, a model from a twin subject (pair 1 twin B individual) and a model inferred from an unrelated subject (pair 2 twin A). The Pgens were highly consistent for the replicate samples: the ratio of the Pgens of the same sequence computed with the two replicate models was in the overwhelming majority of cases below one order of magnitude (Fig. 4A, boxplots). However, for the models inferred from different subjects, both MZ twin and unrelated, the Pgen ratio in some cases reached four or even five orders of magnitude (Fig. 4A). Of note, we did not observe that the ratios for twins were lower than for unrelated individuals: in fact, for some of the antigens (HIV, MERS-CoV), these ratios were higher on average than for the unrelated individuals.

**Figure 4.**
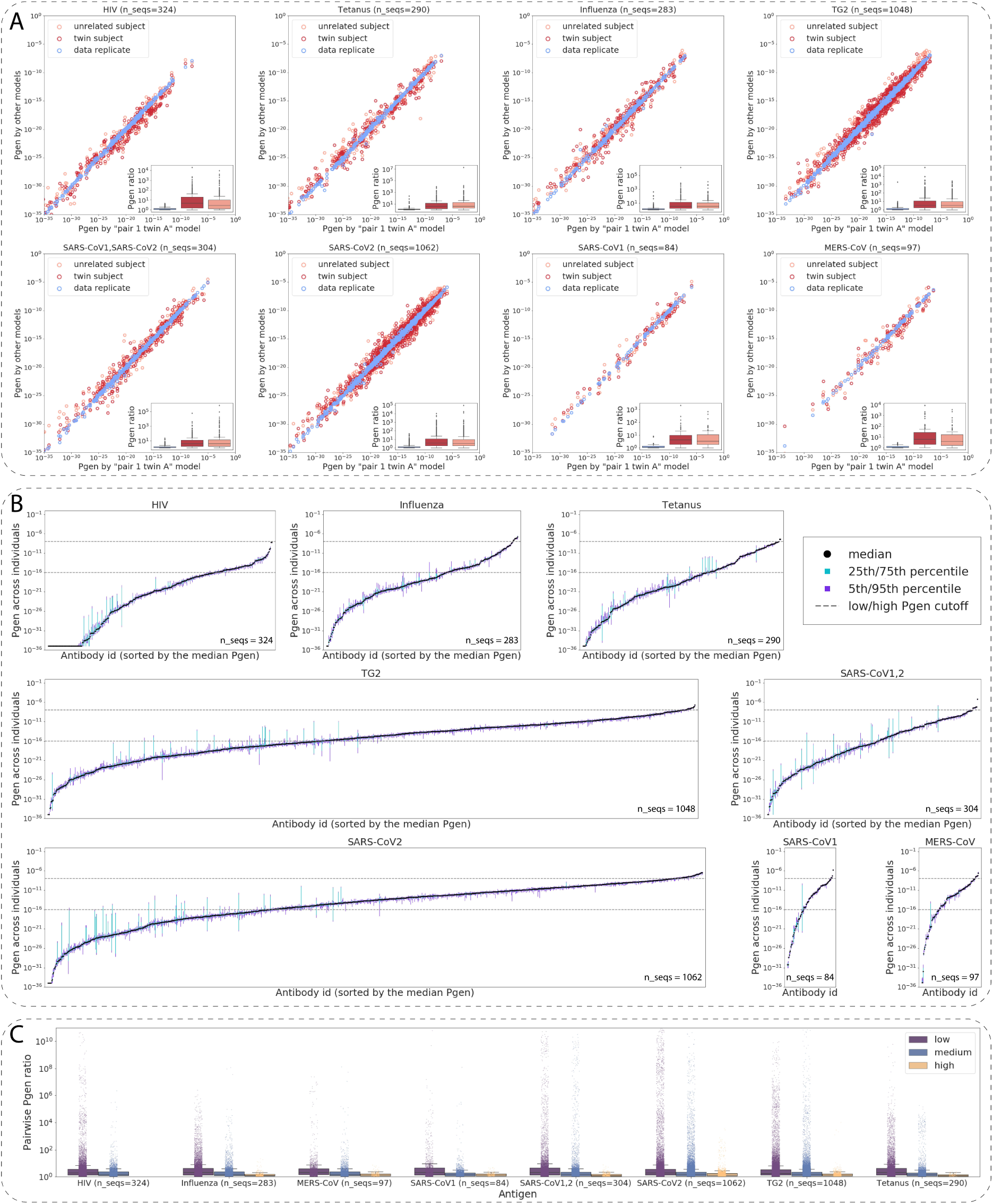
Generation probabilities of antigen-annotated immunoglobulins (CDRH3 sequences) vary by several orders of magnitude within the human population. (A): Pgens of antibody CDRH3 amino acid sequences (annotated with antigen specificity) computed using RGMs corresponding to samples of different levels of immunogenetic similarity: a pair of data replicate models, a pair of models from twin individuals, a pair of models from unrelated individuals. The X-axis always stands for the Pgen as computed with the model corresponding to the pair 1 twin A individual from the HUMAN2 dataset. The Y-axis corresponds to the Pgen as computed with the other model in the pair (data replicate or /twin/unrelated subject). The boxplots show the distribution of the min(x,y)/max(x,y) ratios, i.e., the pairwise difference of Pgens. (B): For each CDRH3 amino acid sequence, we calculated its Pgen as determined by the models corresponding to the 99 individuals from the HUMAN3 dataset. The X-axis itemized each of the CDRH3 sequences tested, the Y-axis denotes the 5th, 25th, 50th, 75th, and 95th percentiles of the 99 Pgens of each CDRH3. (C): Pairwise ratios of the Pgens from (B) by antigen. For each antigen, we divided the CDRH3 amino acid sequences into three groups depending on the sequence’s median Pgen across individuals: low (median Pgen < 10^-16^), medium (10^-16^ ≤ median Pgen < 10^-8^), and high (10^-8^ ≤ median Pgen).

To quantify the Pgen variation of antigen-annotated CDRH3s on a larger scale, we calculated the Pgen of each CDRH3 sequence using the RGM corresponding to the 99 individuals from the HUMAN3 dataset. For each sequence, we calculated the 5th, the 25th, the 50th (median), the 75th, and the 95th percentiles (Fig. 4B). We found that the per-sequence Pgen variation strongly depended on the sequence itself. Variation was especially high for those CDRH3 sequences with mid to low Pgen. To investigate this further, for each CDRH3 sequence, we calculated the pairwise Pgen ratios (Fig. 4C). Then, for each antigen, we split the CDRH3 sequences into three groups according to their median Pgens: “low” group – CDRH3 sequences with median Pgen < 10^-16^ (i.e., sequences that are almost impossible to generate for most individuals. For instance, if a human generates approximately 3×10^13^ throughout their life, then the probability to generate a sequence with Pgen=10^-16^ at least once is approximately 0.003). “Medium” group – CDRH3 sequences with median Pgen between 10^-16^ and 10^-8^, and “high” group – sequences with median Pgen > 10^-8^ (potential public clones: if a sequence can be generated with probability 10^-8^ and the number of human B cells that can be represented in an AIRR-seq sample is approximately 2×10^8^ (Briney et al. 2019), then the sequence will be present in more than 86% of the samples). The variation in the “high” group was lower than in the first two groups: in the “high” group, the variation in most cases stayed within one order of magnitude, while for the “medium” and “low” groups it reached three orders of magnitude for more than a hundred of sequences. Of note, there were no HIV-specific sequences with a median Pgen > 10^-8^; all tested HIV-specific CDR3 sequences belonged to either “low” or “medium” groups.

### Normalized JSD is a sensitive measure to detect subtle repertoire generation parameter differences

Given that the explicit/normalized JSD-based distance depends on the underlying RGMP value distribution, we sought to understand the sensitivity of RGMP parameter inference to variation in the Ig repertoire structure. To this end, we measured the sensitivity of IGoR RGMP inference following small changes in RGMP values. High/low sensitivity means that a given parameter has a great impact/negligible impact on the generated Ig repertoire. This impact can be subsequently measured with the normalized JSD between the initial and the modified RGMP sets.

Starting from an initial RGMP set, we chose a specific RGMP, changed its value, and subsequently used synthetic replicates to test if IGoR was able to correctly determine the shifted value (single-parameter focus, Fig. 5A–D, Fig. S10A-D) and to investigate the sensitivity of the explicit/normalized JSD to such changes (whole-repertoire focus, Fig. 5E–H, Fig. S10E-H). We calculated the explicit JSD between the inferred RGMP sets and the one used for generation (’ground truth RGMP’) and compared it to the explicit JSD across the inferred RGMPs (i.e., between synthetic replicates). For V and J segment (conditional) probabilities, we increased the probability of a segment and then rescaled all other ones accordingly, i.e., we applied a *multiplicative shift* to the chosen segment probability. We chose the RGM inferred from a subsample of 30000 sequencing reads from the pair 1 twin A sample of the HUMAN2 dataset as a starting RGM.

**Figure 5.**
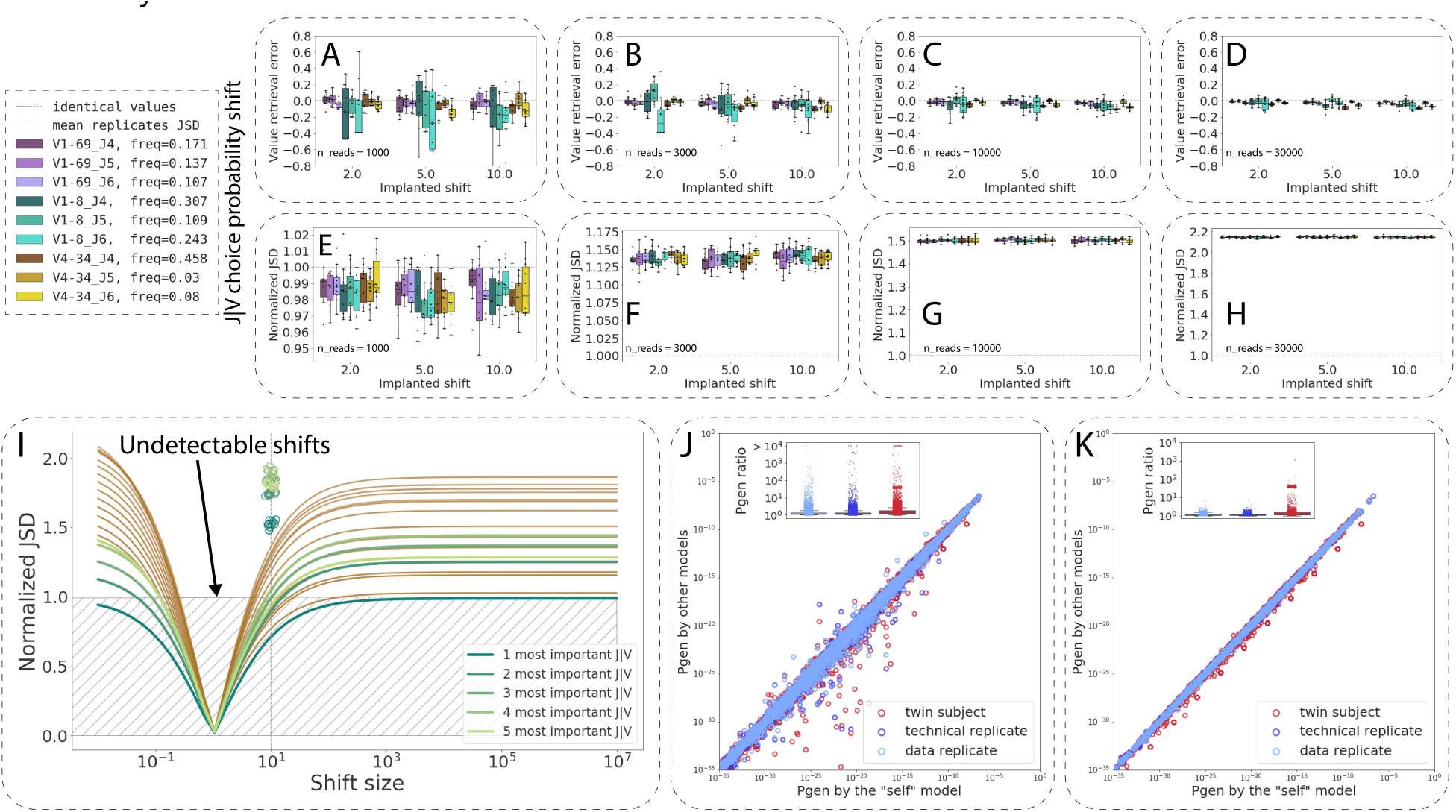
The sensitivity of repertoire generation model parameter inference and their impact on Pgen values vary by repertoire generation model parameter. (A–D): RGMP value retrieval error (the difference between the inferred parameter value and the ground truth one) for RGMPs inferred from synthetic samples that were generated using a modified RGMPs with increased conditional probability to observe a certain J segment given a V segment for synthetic sample sizes of 1000, 3000, 10000, and 30000 sequencing reads (ten synthetic samples for each sample size). The dashed line corresponds to zero difference (i.e., no error observed). (E–H): Normalized JSD between the inferred RGMP sets and the initial modified one (boxes, each box corresponds to the same ten synthetic samples that were used in A–D). Average normalized JSD across the inferred RGMP sets themselves equals one since it is the value used for normalization (i.e. between synthetic replicates, dotted line). (I): All J|V conditional probabilities are ranked by their importance, then the *k* (*k* in [1…20]) most important probabilities are chosen and multiplied by a coefficient from 10^-2^ to 10^6^, the rest is rescaled, to sum up to 1. The X-axis corresponds to this multiplicative coefficient. The Y-axis corresponds to the normalized JSD between the modified model and the unmodified one. The green colors correspond to the first five most important parameters. The circles correspond to the values obtained by generating synthetic samples using the modified model and inferring the parameters back as in E–H. (J) Pgens evaluated on identical sequences using different RGM parameter values (each point corresponds to a single sequence). The X-axis corresponds to the Pgen evaluated using the model parameter values inferred from the same sequences that the Pgens were computed for (the “self” model). The Y-axis corresponds to the Pgens evaluated using other RGMP values (inferred from a data replicate sample, a technical replicate sample, and a sample from a twin subject). The boxplots correspond to the Pgen ratio distributions. (K) Analogous to J but the Pgens were computed only on a set of sequences that consisted of the most impactful combinations of V and J segments (five top pairs as computed in I).

As there are 124 human V segments and tens of thousands of parameters in total (see Methods 3 for a detailed description of model parameters), we restricted our focus to three representative human V gene alleles: a very rare segment (IGHV1-8*01, choice probability of 0.003 in the etalon model), a segment of moderate frequency (IGHV4-34*01, choice probability of 0.024) and a more frequent one (IGHV1-69*06, choice probability of 0.037). We defined the parameter value retrieval error as the difference between the inferred parameter value and the ground truth one used for generating the synthetic data. We calculated the parameter value retrieval error as a function of sample and shift size (Fig. S10A-D, Fig. 5A–D): we used sample sizes of 1000 among 30000 sequencing reads, and we shifted the parameter values multiplicatively by a factor of 2, 5 and 10. We chose these values to not make the modified models too distant from the models inferred from experimental samples: a difference of one order of magnitude in the usage of a given V segment can easily be observed within the population.

For the V segments (IGHV1-8*01, turquoise boxes; IGHV4-34*01, yellow boxes; IGHV1-69*06, violet boxes; Fig. S10), the parameter value estimation was unbiased for IGHV1-8*01 and IGHV4-34*01, and slightly biased for IGHV1-69*06: its probability was systematically underestimated, and the bias was proportional to the multiplicative shift size (i.e., proportional to the parameter value itself). The variance of the parameter value retrieval error decreased with increasing sample size, while the error variance decreased dramatically, as expected for a maximum likelihood estimator like the IGoR inference module. The normalized JSD distance between the inferred RGMP sets and the ground truth one (Fig. S10E-H, Fig. 5E–H) was in most cases higher than the normalized JSD between themselves. This may indicate the presence of a bias in the IGoR synthetic data generation process, i.e., a factor that makes synthetic replicates closer to each other than to the RGMP set used for generation.

For the J segments (IGHJ6, brighter boxes; IGHJ5, medium boxes; IGHJ6, darker boxes, Fig. 5A-D), the parameter value estimation was unbiased, and the variance of the parameter value retrieval error also showed a negative trend with increasing sample size.

To explore the boundaries of IGoR sensitivity, we iteratively modified the RGMPs by shifting twenty of the most important (i.e., of the highest value) parameters and analyzed the normalized JSD between the modified and unmodified model parameters. This time, we used the model inferred from Sample 20 of the MOUSE_NAIVE dataset. All V probabilities (Fig. S10) and J|V (Fig. 5I) conditional probabilities were ranked by their importance, then the *k* (*k* in [1…20]) most important probabilities were chosen and multiplied by a coefficient from 10^-2^ to 10^6^, the other probabilities were rescaled to unity. We then tested the normalized JSD values for the shift size 10 by generating five synthetic samples using the modified model. The normalized JSD values indicate that changing as few as five parameter values is sufficient to significantly change the generated repertoire.

To estimate the impact of the RGMP value variation on the Pgen of a sequence, we computed the Pgens of the same sequences using several different models (Fig. 5J): a model with parameter values inferred from the same sequences that the Pgens were computed for the “self” model (Sample 20, MOUSE_NAIVE), a model inferred from a data replicate sample, a model inferred from a technical replicate sample and a model inferred from a twin subject (Sample 17, MOUSE_NAIVE). This way, the self-model Pgens are more reliable, and we can refer to them as a ground truth.

In order to account for how specific recombination parameters impact the Pgen values, we calculated the distribution of the Pgen ratio between the self-model and the other three (Fig. 5J). The Pgens computed using the replicate models (both the data replicate and the technical one) were closer to the self-model Pgens (the ratio was close to 1 in most cases) than those computed using the twin subject model, where the ratio was below 3 in most cases but reached almost two orders of magnitude for about one percent of the sequences. Interestingly, these high Pgen differences between the twin models remained when we limited our analysis to only those sequences that consisted of the most used V and J segments (top five V-J, i.e., to the sequences with higher Pgens, Fig. 5K). This indicates that the difference in the Pgens is not an artifact that originates from the inference of the low-impactful parameters.

Collectively, our data support the view that certain parameters impact IGoR RGMP inference and Pgen evaluation to a substantially larger extent than others and that an artificial perturbation of as few as five parameter values is sufficient to produce an observable difference in the generated repertoire. Moreover, our analysis supports the view that the JSD is a sufficiently sensitive measure to detect this difference.

## Discussion

We demonstrated that Ig VDJ recombination rules (RGM) and, consequently, Ig sequence generation probabilities differ across individuals, even between homozygous human twins and inbred mice (Fig. 1). Our approach (which we call desYgnator) relies on a hierarchy of experimental controls as well as information and statistical theory and provides new recommendations for unbiased comparison of RGM between individuals, that were previously thought to be identical (Fig. 2). Our results indicate that inter-individual differences in VDJ recombination are not only due to genetic differences in germline gene repertoires (e.g., germline gene polymorphisms, structural variation) but also due to non-genetic differences (e.g., epigenetics). Indeed, it has been previously shown that epigenetic mechanisms intervene in the regulation of VDJ recombination (Pulivarthy et al. 2016). Specifically, our work shows that individuals do not only differ in their recombined expressed repertoire (Dupic et al. 2020) but that already the individual sources (RGM) of each expressed repertoire differ. We found that the distance between RGM of twin subjects is on average 5% lower than between RGM of unrelated subjects (Fig. S9). However, this does not hold when the distances are measured for V-segment-agnostic RGM (Fig. S5), suggesting that non-genetic factors account for approximately 90% of the RGMP difference in general and for almost all on the V-segment-unrelated RGMP. Thus, our results suggest that V-segment-unrelated RGMP are not genetically heritable (Glanville et al. 2011). Specifically, we posit that antigen binding governed by the V-region (e.g., influenza, Avnir et al. 2016) is partially genetically heritable (Venkataraman et al. 2021) whereas cases where most of the binding is governed by the V-agnostic portion (i.e., CDRH3), is not genetically heritable. Of note, in the largest structural antibody-antigen binding data set to date, we previously showed that the CDRH3 is the sole obligate region for antibody binding (Akbar et al. 2021). Therefore, our analysis suggests that for a large number of antigens, Ig-driven antigen recognition is not genetically heritable.

We found that, for unrelated subjects, the aforementioned RGMP difference correlates with the number of differing IGHV alleles (Fig. 3C). Surprisingly, the correlation can even be observed with V-segment-agnostic RGM (Fig. S8C). We speculate that this may be caused by at least two reasons: (i) individuals that are more different in the IGHV locus are more genetically dissimilar in general, hence they are more likely to have polymorphisms in other parts of the IG superlocus or to differ in non-genetic factors that affect the VDJ recombination. (ii) Different sets of IGHV germline segments inherently affect the inference of other RGMP: the model has to explain the data with other VDJ scenarios.

The study of AIR sequence enrichment, for instance, to identify antigen-expanded sequences, previously assumed a unique RGM shared by all individuals of a species (Marcou et al. 2018). We showed one effect of the variation in RGMPs by applying different models to a given set of CDRH3 sequences (Fig. 4, Fig. 5). We found that the extent of correlation between the Pgens was consistent with the degree of immunogenic similarity between the models, i.e., the correlation was higher for the models inferred from replicate samples than from samples obtained from different subjects. We also found that the Pgen of the same Ig sequence can differ by several orders of magnitude between individuals, and, consequently, that the effectively available Ig sequence space varies from one individual to another. Thus, future methods for identifying antigen-specific sequences may require considering individual-specific RGM.

In our work, we considered several types of replicate samples: biological replicates (DeWitt et al. 2016), technical replicates (Greiff et al. 2017a), and *in silico*-constructed data replicates and synthetic replicates. Despite providing a reliable baseline for estimation of the RGMP variation when inferring from samples from the same subject, these types of replicates do not span all steps of AIRR-seq sample generation. For example, our analysis reveals the importance of quantifying the extent to which RGMP differs across different library preparation protocols, e.g., RACE, multiplex, influence of UMI usage (Barennes et al. 2021; Vázquez Bernat et al. 2019; Menzel et al. 2014; Khan et al. 2016).

Our approach is highly sensitive to subtle RGMP modifications: we showed that an artificial perturbation of as few as five parameter values can be detected using desYgnator (Fig. 5).

In addition to the biological findings and the statistical framework developed, we provide guidelines for RGM analyses. Specifically, we find that more than a minimum number of sequencing reads (10,000) are necessary for the faithful inference of RGMP (Fig. 5). In contrast to current practice, we show that RGMP datasets significantly differ from one another and therefore suggest inferring RGMPs for each individual, especially when an increased signal-to-noise ratio is desired. Furthermore, it is commonly held that RGMPs need to be either inferred from out-of-frame sequences or very early B/T-cell stages (Marcou et al. 2018). While we agree that B/T-cells unaffected by any selection (e.g., pro-B cells) would represent the ideal data for RGMP inference here, we inferred RGMPs from both in- and out-of-frame sequences of either pre-B or naïve B cell repertoires (see Methods 4 for details and reasoning). We also provide a guideline for preprocessing human AIRR-seq data with respect to the correct set of germline alleles for each individual (Fig. 6, Fig. S1). We did not use previous methodology developed for novel allele inference (Corcoran et al. 2016; Gadala-Maria et al. 2019; Ralph and Matsen 2016; Zhang et al. 2016) because our goal was to provide stable input for the RGMP inference step and, after that, to compare RGMP across individuals which required a general database of validated alleles. We, therefore, discarded potentially novel alleles in favor of previously validated ones.

**Figure 6.**
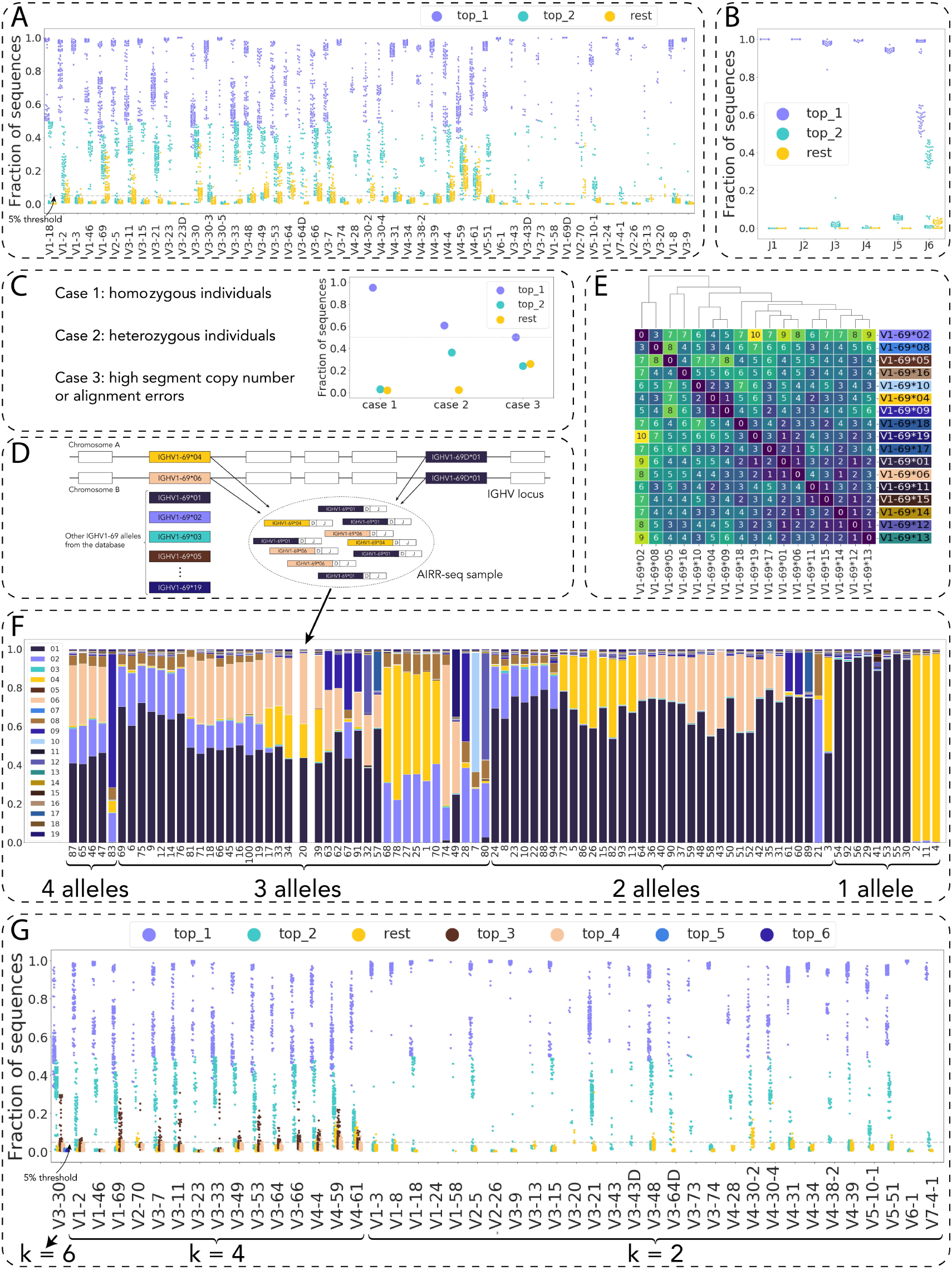
Impact of individualized restriction of germline gene database on allelic assignment in AIRR-seq samples. (A) The proportion of the two most frequent alleles for 100 individuals from the HUMAN3 dataset. Each column on the plot corresponds to one IGHV gene. For each sample (individual), three data points are shown: the fraction of sequencing reads assigned to the two most frequent alleles (violet: top_1, turquoise: top_2) and the total fraction of all remaining alleles (yellow: rest). The dashed gray line corresponds to the 5% threshold above which we consider allele fractions as not due to sequencing/PCR errors. Alignments were performed using MiXCR v3.0.12. (B) Analogous to A but for IGHJ genes. (C) Per-gene alignment scenarios can be divided into three general cases. Case 1: a high fraction (> 90%) of sequencing reads assigned to the most frequent allele, and negligible fractions (< 5%) of sequencing reads assigned to the remaining ones (which suggests a homozygous individual for a given gene). Case 2: comparable fractions of sequencing reads assigned to the first and the second most frequent alleles and a low (< 5%) total fraction of the remaining ones (suggesting a heterozygous individual). Case 3: a high (> 5%) total fraction of the remaining alleles, suggesting a systematic alignment bias likely caused by gene duplication, which may lead to the presence of three or more alleles of the current gene. (D) Illustration of the generation of an AIRR-seq sample that contains more than two alleles of the same gene. (E) Example: the sequence diversity of IGHV1-69 alleles as quantified by the edit (Levenshtein) distance. (F) Example: the proportion of the IGHV1-69 alleles across the 99 individuals of the HUMAN3 dataset: one column corresponds to one individual. The columns are ordered by the number of alleles present, and within each category for allelic diversity (4 alleles, 3 alleles, etc.) by the most frequent alleles. (G) The fraction of the remaining alleles is low and is below the 5% threshold for sequencing and PCR error, for most of the genes and most of the individuals after performing the full pipeline, i.e., with realignment the sequences against the restricted set of alleles (see Methods 3 and Fig. S1).

Machine learning is increasingly used for AIRR classification both on the sequence (Akbar et al. 2021; Friedensohn et al. 2020; Isacchini et al. 2021; Greiff et al. 2017b) and repertoire-level (Shemesh et al. 2021; Emerson et al. 2017; Sidhom et al. 2021; Pavlović et al. 2021). Future studies will need to investigate whether differences in RGM also impact repertoire classification (Greiff et al. 2020). If epigenetic factors influence AIRR architecture, it will be interesting to investigate whether aging or shorter biological rhythms change the rules of VDJ recombination over time. This may be done via analyzing longitudinal data (Mitsunaga and Snyder 2020), or pre- and post-puberty data from monozygotic twins (as epigenetic differences have been shown to arise after puberty, Fraga et al. 2005). Another way to investigate the non-heritable factors that impact VDJ recombination is to study samples from different cell populations in the same individual (from pro- to naïve and even memory B cells), as it will allow quantifying the influence of the negative and positive selection (Nemazee 2017; Robert et al. 2021).

It is important to mention that although we limited our attention to B-cell datasets in this study, our approach is directly applicable to T-cell receptor repertoires, provided the availability of the appropriate high-quality data – RNA sequences of non-antigen-experienced T cells from biological and/or technical replicates, as well as from monozygotic twins and unrelated subjects.

In the future, it will be interesting to study whether individual differences in RGMP lead to differences in the propensity to generate antigen-specific (for instance auto-reactive) sequences (Shemesh et al. 2021) and, consequently, to the existence of individualized holes in the repertoire (Perelson and Oster 1979). These analyses will require large-scale naïve (unselected) and disease-linked AIRR-seq data (Watson, Glanville, and Marasco 2017). Furthermore, a further analysis of the link between germline polymorphisms and RGMP based on population-wide genomic data that include non-coding regions (Mikocziova et al. 2020) will also be of interest. Another avenue to extending our analysis is to account for unconventional cases of VDJ recombination, such as the absence of D segments in TCR*β* chains (Greef and Boer 2021) or the occurrence of multiple D segments in IGH chains – VDDJ recombination (Safonova and Pevzner 2020).

Our work has implications for vaccine development. There is an increasing interest in understanding whether B cells that are to be targeted by immunogens exist within the naïve B cell repertoire of most people in a population of interest (“public clones”, Greiff et al. 2017; Elhanati et al. 2018), and if those B cells occur at a high enough precursor frequency such that they have a high likelihood to become activated in response to immunizations. These considerations relate to both V gene usage but also germline gene polymorphisms (Lee et al. 2021; Sangesland et al. 2019). Here we show that one factor towards the variation in naïve B cell precursor frequencies and repertoires is personalized VDJ recombination models. Nowadays, high-throughput sequencing technologies allow deep sequencing of naive Ig repertoires, thus enabling the integration of VDJ recombination models into iterative individualized immunogen design pipelines to advance vaccine discovery (Lee et al. 2021).

## Methods

### Methods 1. Experimental immunoglobulin sequencing data

We analyzed four publicly available Ig experimental datasets:

<u>MOUSE_PRE</u> from Greiff et al. 2017 (ArrayExpress E-MTAB-5349): IgH pre-B cell samples obtained from nineteen inbred SPF C57BL/6 mice, sequenced on the Illumina MiSeq platform (2×300 bp). For one of the mice, Greiff and colleagues prepared two technical replicates, resulting in twenty samples in total.

<u>MOUSE_NAIVE</u> from Greiff et al. 2017 (ArrayExpress E-MTAB-5349): IgH naïve B cell samples obtained from twenty inbred SPF C57BL/6 mice, sequenced on the Illumina MiSeq platform (2×300 bp). For one of the mice, Greiff and colleagues prepared two technical replicates, resulting in twenty-one samples in total.

<u>HUMAN1</u> from DeWitt et al. 2016 (Dryad Digital Repository doi:10.5061/dryad.35ks2): IgH naïve B cell samples obtained from three 25–40-year-old Caucasian male donors, sequenced on the Illumina HiSeq platform (1x130 bp spanning CDR3). For one of the donors, DeWitt and colleagues prepared two biological replicates, resulting in four samples in total.

<u>HUMAN2</u> from Rubelt et al. 2016 (Sequence Read Archive SRP065626): IgH naïve B cell samples obtained from five pairs of adult monozygotic twins (ten samples in total), sequenced on the Illumina MiSeq platform (2×300 bp).

<u>HUMAN3</u> from Gidoni et al. 2019 (European Nucleotide Archive PRJEB26509): IgH naïve B cell samples obtained from one hundred individuals from Norway; 48 healthy controls (out of which 28 blood bank donors and 20 healthy individuals), and 52 patients with celiac disease; sequenced on the Illumina MiSeq platform (2×300 bp). We discarded one of the samples (S97) due to the poor quality of the sequencing reads.

For further experimental details on each dataset, please refer to the respective publications.

### Methods 2. An approach to building personalized repertoire generation models that are robust to allelic variability of IGHV genes

Evidence published over the last couple of years has demonstrated an extensive polymorphic and structural diversity in the immunoglobulin germline locus (Watson et al. 2017; Lees et al. 2020; Collins et al. 2020; Yu et al. 2017; Martins et al. 2021; Khatri et al. 2020; Bernat et al. 2021; Mikocziova et al. 2021). These findings indicate that the RGMs used to annotate each individual’s immune receptor sequences with accurate Pgens need to be built for each individual separately with the individualized set of alleles. The human IGHV locus is highly diverse (Watson and Breden 2012a; Watson et al. 2017) and there exist multiple alleles for the majority of IGHV genes in the IMGT and OGRDB databases, e.g., up to 19 alleles for genes IGHV1-69 and IGHV3-30) (Lefranc 2001; Lees et al. 2020). AIRR-seq VDJ gene annotation tools (e.g., MiXCR by Bolotin et al. 2015; IgBLAST by Ye et al. 2013; the annotation module of IGoR by Marcou, Mora, and Walczak 2018; partis by Ralph and Matsen 2016) rely solely on a given set of germline genes and alleles from a chosen germline gene database. Any sequencing read within a sample needs to be aligned to all alleles in a germline gene database. Taking a full germline database as reference behavior might be problematic for the inference of personalized RGMP since each individual may only possess a subset of the alleles present in the germline database.

Theoretically, one human individual cannot have more than two alleles of the same gene. However, the structure of the IGHV locus allows for exceptions to the “two alleles per gene” rule (Mikocziova et al. 2020; Ford et al. 2020) due to the existence of repeated segments. Additionally, sequencing and PCR errors may lead to misalignments (we set the upper bound for the fraction of erroneous allele assignments due to sequencing/PCR errors to 5%, see Fig. S4).

For each human individual of each dataset in this study (HUMAN1, HUMAN2, and HUMAN3 datasets, see Methods 1), we aimed to construct an individually restricted germline gene database that contained only those validated alleles that are present in the given individual. This restricted database is thus a subset of all available validated alleles present in IMGT and OGRDB.

In this section, we show an example of deriving such an individually restricting process via analyzing the HUMAN3 dataset, as it is the most diverse one (99 unrelated individuals). The pipeline itself is described in the following.

For each IGHV (Fig. 6A) and IGHJ (Fig. 6B) gene and each individual, we calculated the fraction of sequencing reads (see Methods 4 for the definition of a sequencing read) assigned to the two most frequent alleles (top_1 and top_2), as well as the fraction of sequencing reads assigned to all remaining alleles (rest). This calculation exposed three genomic scenarios (Fig. 6C):

<u>Case 1</u>: a very high fraction of reads (> 90%) assigned to the most frequent allele and negligible fractions of reads assigned to the lower frequency alleles. We assumed this scenario corresponds to an individual that is homozygous for this gene. This case was found for at least four IGHV genes for 97 out of 99 individuals and for IGHJ1-IGHJ5 genes for all individuals. IGHJ6 gene belonged to this case for 44 out of 99 individuals.

<u>Case 2</u>: non-negligible fractions (> 5%) of sequencing reads assigned to the first and the second most frequent alleles and a very low rest-fraction (< 5%) – suggesting a heterozygous individual. This case was found for at least one IGHV gene for 93 out of 99 individuals. The only IGHJ gene this case was found for was IGHJ6 (for 54 out of 99 individuals).

<u>Case 3</u>: a high rest-fraction (> 5%) suggesting that this individual has more than two alleles of the current gene (excluding errors leading to a systematic misalignment). This case was found for at least one IGHV gene for 93 individuals. No IGHJ gene belonged to Case 3 (Fig. 6B), which may be explained by the fact that the IGHJ locus is composed of fewer genes and fewer alleles per gene (Watson and Breden 2012a; Gidoni et al. 2019; Peres et al. 2019).

To illustrate Case 3, we visualized the allelic complexity of the IGHV1-69 gene (Fig. 6D–F), known to be crucial for generating potent neutralizing antibodies (Avnir et al. 2016; Brouwer et al. 2020). Specifically, in individual #20 of the HUMAN3 dataset, sequencing reads were aligned to eight alleles of IGHV1-69 (gene 69 in IGHV gene family 1). IGHV1-69 has a known duplication in the IMGT database named IGHV1-69D (Giudicelli et al. 2005). The duplicated gene has one allele (IGHV1-69D*01), which it shares with the original gene (see Fig. 6D). It is not feasible to determine the origin of the shared allele from a recombined VDJ sequence, so it is *de facto* possible to observe three different alleles of IGHV1-69: two originating from the gene itself and the third one from the duplication. In the aforementioned individual 20, only three alleles out of eight had above-the-threshold fractions of the sequencing reads annotated with IGHV1-69: IGHV1-69*01, IGHV1-69*04, and IGHV1-69*06, which is exactly the case described in the previous sentence: IGHV1-69*04 and IGHV1-69*06 might have originated from the gene IGHV1-69 while IGHV1-69*01 might be, in fact, IGHV1-69D*01 having originated from a duplication.

IGHV1-69 has 19 alleles of this gene, some of which are very close to each other, i.e., differ by a single nucleotide substitution (Fig. 6E). There were five individuals (namely, individuals 46, 47, 65, 83, 87) that had four alleles (Fig. 6F) with fractions above the 5% threshold (i.e., each of these alleles accounted for more than 5% of the sequencing reads assigned to IGHV1-69), thirty-seven individuals with three frequent alleles, and thirty-four individuals with two frequent alleles (which can signify a normal heterozygous gene or a homozygous with a mutated copy). The remaining individuals had a single IGHV1-69 allele with a frequency above the threshold. To summarize, human AIRR-seq samples *can* have more than two alleles of the same IGHV gene, and one should treat these cases separately to avoid systematic biases in the fitted RGMs – or to ensure that these systematic biases are the same for all considered AIRR-seq samples, which would allow comparing the RGMP inferred from these samples.

Finally, we individually restricted germline gene databases as follows (see Methods 3): we set the maximum possible number of alleles (which we denoted by *k*) for each gene individually (Table 1) based on the information on its potential copy number, duplications in the databases and the high rest-score from Fig. 6A, and trimmed all alleles after the 2nd-CYS, amino acid position 104 in the IMGT unique numbering scheme (Lefranc et al. 2003). Then, we determined the *k* most frequent alleles of the given gene in the sample in order to restrict the RGM to these alleles only.

**Table 1.**
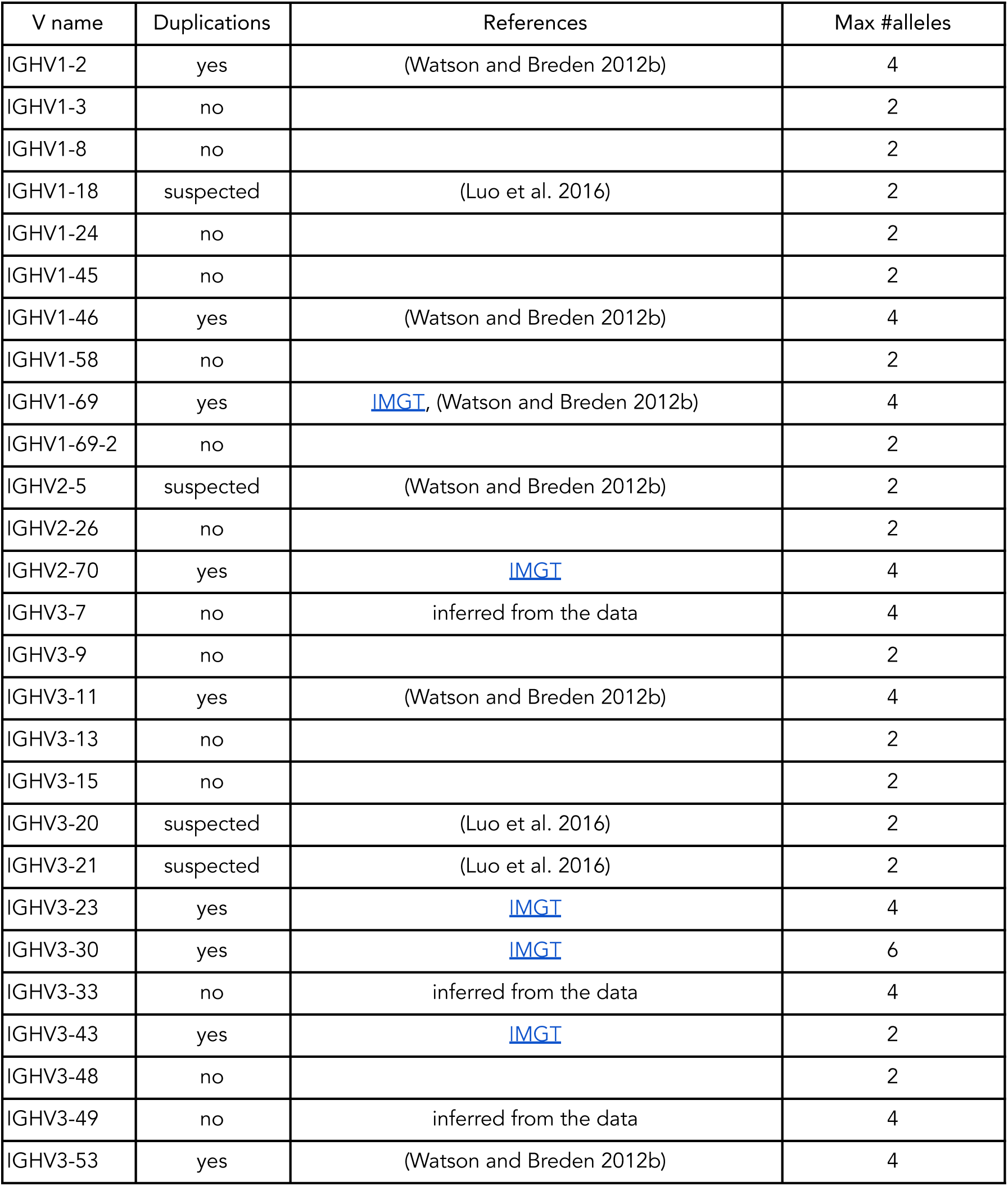
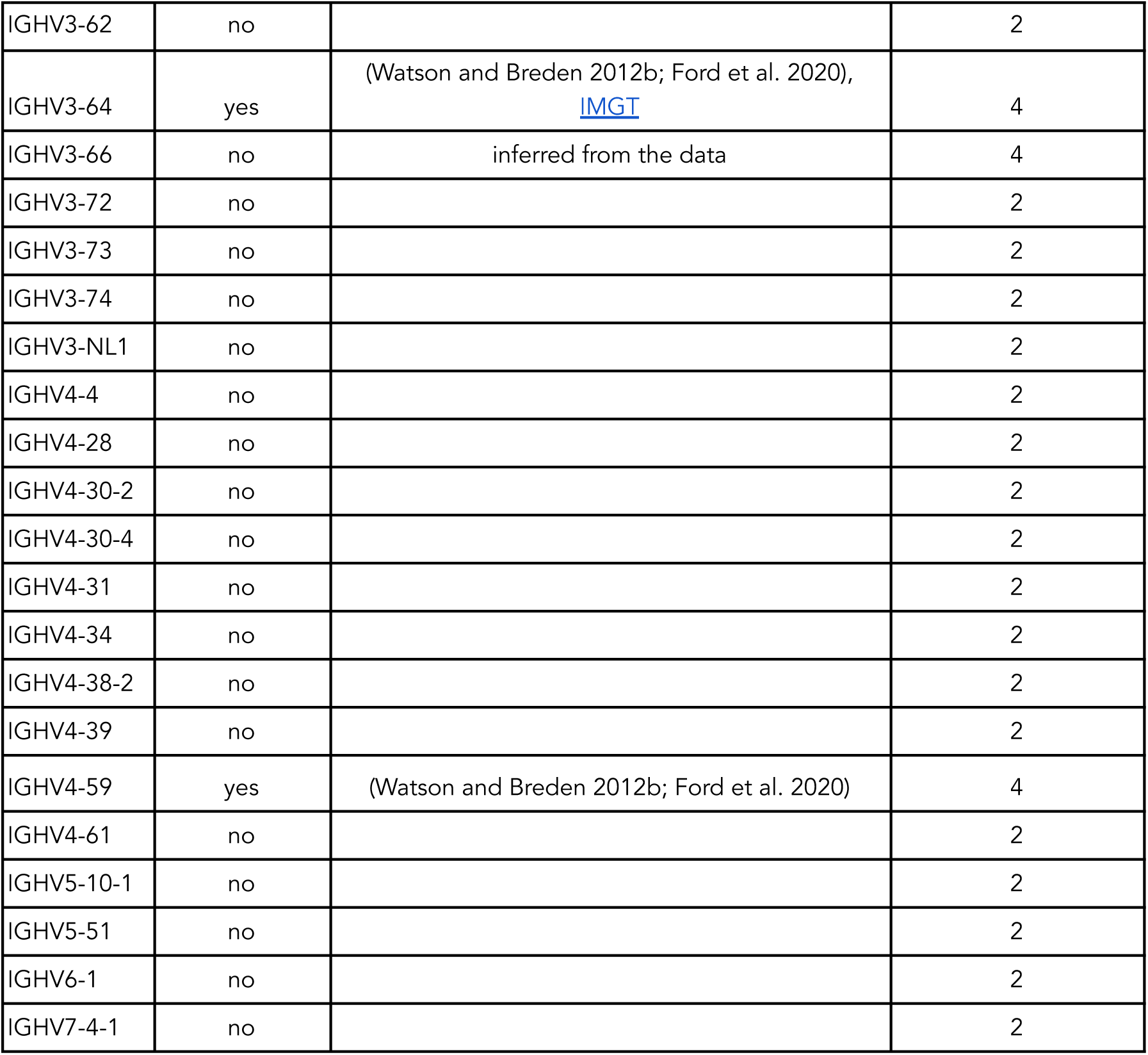
(relates to Fig. 6). Maximum number of alleles set for each IGHV gene. These numbers are based on the previously known information about IGHV copy numbers and on the high fractions of remaining alleles (not top_1 or top_2 alleles in terms of frequency) in the HUMAN3 dataset (Fig. 6A).

| V name | Duplications | References | Max #alleles |
| --- | --- | --- | --- |
| IGHV1-2 | yes | (Watson and Breden 2012b) | 4 |
| IGHV1-3 | no |  | 2 |
| IGHV1-8 | no |  | 2 |
| IGHV1-18 | suspected | (Luo et al. 2016) | 2 |
| IGHV1-24 | no |  | 2 |
| IGHV1-45 | no |  | 2 |
| IGHV1-46 | yes | (Watson and Breden 2012b) | 4 |
| IGHV1-58 | no |  | 2 |
| IGHV1-69 | yes | IMGT, (Watson and Breden 2012b) | 4 |
| IGHV1-69-2 | no |  | 2 |
| IGHV2-5 | suspected | (Watson and Breden 2012b) | 2 |
| IGHV2-26 | no |  | 2 |
| IGHV2-70 | yes | IMGT | 4 |
| IGHV3-7 | no | inferred from the data | 4 |
| IGHV3-9 | no |  | 2 |
| IGHV3-11 | yes | (Watson and Breden 2012b) | 4 |
| IGHV3-13 | no |  | 2 |
| IGHV3-15 | no |  | 2 |
| IGHV3-20 | suspected | (Luo et al. 2016) | 2 |
| IGHV3-21 | suspected | (Luo et al. 2016) | 2 |
| IGHV3-23 | yes | IMGT | 4 |
| IGHV3-30 | yes | IMGT | 6 |
| IGHV3-33 | no | inferred from the data | 4 |
| IGHV3-43 | yes | IMGT | 2 |
| IGHV3-48 | no |  | 2 |
| IGHV3-49 | no | inferred from the data | 4 |
| IGHV3-53 | yes | (Watson and Breden 2012b) | 4 |
| IGHV3-62 | no |  | 2 |
| IGHV3-64 | yes | (Watson and Breden 2012b; Ford et al. 2020),<br>IMGT | 4 |
| IGHV3-66 | no | inferred from the data | 4 |
| IGHV3-72 | no |  | 2 |
| IGHV3-73 | no |  | 2 |
| IGHV3-74 | no |  | 2 |
| IGHV3-NL1 | no |  | 2 |
| IGHV4-4 | no |  | 2 |
| IGHV4-28 | no |  | 2 |
| IGHV4-30-2 | no |  | 2 |
| IGHV4-30-4 | no |  | 2 |
| IGHV4-31 | no |  | 2 |
| IGHV4-34 | no |  | 2 |
| IGHV4-38-2 | no |  | 2 |
| IGHV4-39 | no |  | 2 |
| IGHV4-59 | yes | (Watson and Breden 2012b; Ford et al. 2020) | 4 |
| IGHV4-61 | no |  | 2 |
| IGHV5-10-1 | no |  | 2 |
| IGHV5-51 | no |  | 2 |
| IGHV6-1 | no |  | 2 |
| IGHV7-4-1 | no |  | 2 |

To visualize the effect of this preprocessing pipeline on the fractions of the remaining alleles (i.e., not the 2 or the *k* most frequent ones), we recomputed the fractions from Fig. 6A after trimming the alleles and setting the per-gene *k* values but before restricting the set of alleles to the *k* most frequent ones (Fig. 6G). This procedure led to non-zero rest-scores <5% in the majority of cases suggesting that we successfully eliminated (most of) the systematic alignment bias. The last step of the pipeline consisted of restricting the allele database and realigning the sequencing reads, which eliminated rest score assignments (Fig. S3). This preprocessed dataset was used for all Pgen inferences.

It is important to mention that, when we applied the described workflow to the HUMAN2 dataset, the individually restricted allele databases of monozygotic twins were not identical due to the presence of weakly expressed alleles. However, the allele sets of monozygotic twins were more similar than those of unrelated individuals.

### Methods 3. A detailed description of the pipeline for building individually restricted germline allele databases

Diploidy and structural variations in the IGHV locus, along with the high similarity between the IGHV segments render annotation of AIRR-seq data a complex problem. The majority of the currently available annotation tools (Bolotin et al. 2015; Ye et al. 2013; Brochet et al. 2008; Ralph and Matsen 2016) may suggest several V segments for each sequencing read. Each sequencing read is processed independently, which may result in the presence of any number of alleles per gene in the annotations for a given sample. Thus, there is a need (see Fig. 6) for a method to accurately restrict the set of IGHV alleles in AIRR-seq data for each individual. To address this need, we employed the following strategy (Fig. S1): <u>Step 1</u>: we constructed a library of V allele nucleotide sequences based on IMGT (Lefranc 2001, downloaded on 2020-05-29) and OGRDB (Lees et al. 2020) downloaded on 2020-05-29), databases, and trimmed all sequences after the second Cysteine, nt position 312 according to the IMGT unique numbering scheme (Lefranc et al. 2003). We performed trimming to ensure the absence of alignment errors which may be caused by nucleotides located after position 312, i.e., in the VD junction (this problem was also noted by (Mikocziova et al. 2020)). <u>Step 2</u>: for those alleles of the same gene that became identical after trimming (namely, IGHV1-46*01=*03, IGHV3-7*02=*04, IGHV3-30*03=*18, IGHV3-33*01=*06, IGHV3-66*01=*04, IGHV4-28*01=*03), we deleted all but one copy. If there were different genes that shared identical alleles (such as IGHV2-70*04 and IGHV2-70D*04), we considered these genes operationally indistinguishable (Luo et al. 2019) and combined their allele sets (again, leaving only one copy of the duplicated allele). <u>Step 3</u>: for each gene, we set a number *k* – the maximum number of alleles that we expect to observe in a sample: default, *k* = 2, for diploid species. However, for some individuals, there may exist more than two alleles for those genes whose allele sets were combined on the previous step (Fig. 6D). For those genes, we set k = 4 (and we set k = 6 for IGHV3-30 as its allele set originates from 3 genes: IGHV3-30, IGHV3-30-3 and IGHV3-30-5). We also set k = 4 for those genes that are known to have multiple copies (Watson and Breden 2012a; Ford et al. 2020; Luo et al. 2019), see Table 1), <u>Step 4</u>: with the modified allele library, we annotated the AIR nucleotide sequencing reads of the given sample using MiXCR (version 3.0.12). <u>Step 5</u>: for each gene, we detected the *k* most frequent alleles, restricted the allele library to these alleles, thus obtaining an individually restricted allele library. <u>Step 6</u> consisted of annotating all AIR sequencing reads again using the individually restricted allele library. Note that the blue path on Fig. S1 (steps 1–3) corresponds to preprocessing the allele library and hence those steps need to be performed only once if the original allele database has not been updated. The orange (steps 4–5, computing the individually restricted allele library) and violet (step 6, annotating the sequencing reads) paths, however, must be repeated for each AIRR-seq sample.

For murine data, we only cleaned the sequencing reads (i.e., assembled the contigs) using MiXCR (Bolotin et al. 2015): we used the MiXCR-computed clones, each clone multiplied by its abundance. We did not apply the allele preprocessing workflow, since all considered subjects were inbred mice, so by definition, they share identical germline gene sets.

Of note, when aligning the sequences with IGoR for the subsequent model inference, we used an increased D gene alignment score threshold Fig. S6): with the default threshold, the alignment module of IGoR produced a huge amount of alignments that did not significantly impact nor the fitted model parameter values nor the sequence Pgens evaluated using this model.

### Methods 4. On the strategy not to filter unproductive sequences

To reduce the amount of noise caused by PCR amplification, we assembled the clone contigs using MiXCR (Bolotin et al. 2015). So as not to lose the information contained in the clone frequencies, we multiplied each clone by the number of reads used to construct it. We refer to these multiplied clone nucleotide sequences as “sequencing reads”.

Although Marcou and colleagues suggest (Marcou et al. 2018) to filter out productive sequences for IGoR inference, we inferred RGM from both productive and unproductive reads. The main reason for this strategy is that we intended to fit the IGoR model to the most general recombination rules: if the VDJ recombination can produce a sequence (productive or unproductive), then we should take this sequence into consideration. Consequently, since we did not separate productive sequences from unproductive ones anywhere, we did not introduce a definition of unproductive sequences. Furthermore, we only worked with data from flow-cytometry sorted non-antigen-experienced cells (pre- and naïve B cells). As hypothesized in the studies in the field (Marcou et al. 2018; Sethna et al. 2020; Desponds et al. 2021), we argue that, although those cells have already undergone selection, the bias introduced by the selection is negligible when compared to the inter-individual differences: if selection impact on RGMP was too high, we would have seen that RGMP sets inferred from naïve B cell samples obtained from different subjects were indistinguishable (i.e., the distance would have been comparable to the distance caused by the noise). However, we see the opposite: although RGMP sets inferred from naïve B cell samples were closer to each other than RGMP sets inferred from pre-B cell samples, they were still clearly distinct (Fig. 2E–G).

### Methods 5. Using Jensen-Shannon divergence to compare repertoire generation model parameters

Kullback-Leibler divergence (Kullback and Leibler 1951) measures the difference between two probability distributions *P* and *Q*. For discrete probability distributions defined on the same probability space, the Kullback-Leibler divergence is defined as 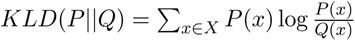. Marcou and colleagues calculated the Kullback-Leibler divergence (KLD) of the IGoR-estimated model parameter values to the ground truth ones to validate the IGoR inference module (Marcou, Mora, and Walczak 2018). We argue that a similar method can be used for comparing IGoR models inferred from different AIRR-seq samples. We used the Jensen-Shannon divergence, which is a smoothed symmetric version of the Kullback-Leibler divergence: 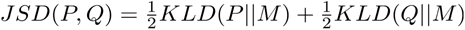, where 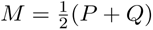.

Jensen-Shannon divergence is symmetric, non-negative and due to the smoothing, its square root satisfies the triangle inequality. Thus, the square root of the JSD can be used as a distance between IGoR models – and between AIRR-seq samples the models were inferred from.

The Kullback-Leibler divergence (and, hence, the Jensen-Shannon divergence) between two multivariate distributions can be decomposed into a sum of several components if components of the distributions are conditionally independent. Applied to the IGoR RGM, this means that the JSD (‘explicit JSD’) between two IGoR models can be decomposed into seven additive terms. We used the default set of IGoR model parameters for IgH:

- V choice probabilities (a vector of length equal to the number of V segments in the database, #V). For the human model, we considered 124 V genes. For the mouse model, we considered all 238 V gene alleles.
- conditional J | V choice probabilities (a matrix of size #V ✕ #J; 6 J genes for the human model, 4 for the mouse model).
- conditional D | V,J choice probabilities (a three-dimensional tensor of size #V ✕ #J ✕ #D; 35 D genes for the human model, 25 for the mouse model)
- conditional delV | V deletion profiles (a matrix of size #V ✕ maximum deletion length=41)
- conditional delJ | J deletion profiles (a matrix of size #J ✕ maximum deletion length=41)
- conditional delD3’ | D and delD5’ | D,del_D3’ deletion profiles (a matrix of size #D ✕ maximum deletion length and a three-dimensional tensor of size #D ✕ maximum deletion length ✕ maximum deletion length=41)
- VD insertion profiles (a vector of length maximum insertion length=41)
- DJ insertion profiles (a vector of length maximum insertion length=41)

The explicit JSD between two sets of IGoR parameter models *P*_1_ and *P*_2_ can be then written as the sum of the JSDs of the joint probabilities of conditionally independent recombination events:

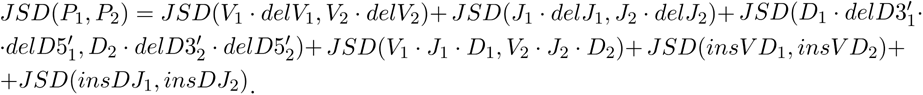

We computed a lower bound for the variation of the explicit JSD introduced by the noise by recreating this variation in the most controlled setting: for each dataset, for a given sample size of N sequencing reads (N in [1000, 3000, 10000, 30000]), we used an IGoR model M_0 (inferred from the first experimental sample of each dataset), generated ten pairs of synthetic samples of N sequencing reads each and computed the explicit JSD of the IGoR-inferred RGMP values from these sequencing reads, within the pairs (synthetic replicates on Fig. 2A). Thus, we obtained a set of values of explicit JSD between datasets generated using *the same theoretical RGMP*. To obtain the normalized JSD between two RGMP sets, we divide their explicit JSD by the mean of the explicit JSDs obtained using generated synthetic samples. We used the same pre-computed synthetic samples within one species: if the normalized JSD for RGMP sets inferred from samples *E*_1_ and *E*_2_ is to be calculated, we set *normalizedJSD*(*E*_1_, *E*_2_) = *explicitJSD*(*E*_1_, *E*_2_)/*mean_ij_explicitJSD*(*S_i_*,*S_j_*) where the same synthetic replicates *S_i_* are used for any samples for a given species (human/mouse).

For the human data, we applied the allele preprocessing pipeline (Fig. S3) from Methods 3 to remove the systematic bias in the model inference. Subsequently, we summed the parameters corresponding to alleles of the same gene and compared different subjects on the gene, not on allele level. We did not apply the allele preprocessing workflow for the murine data, since all subjects share by definition (inbred mice) identical germline gene sets.

In this study, we limited the maximum sample size to 30000 sequencing reads due to the high time and memory consumption by IGoR (inferring RGMP values from all considered samples took about 300000 CPU hours).

### Methods 6. A statistical test for comparing repertoire generation models

To test if the RGMP sets for two AIRR-seq samples A (containing N sequencing reads) and B (containing K sequencing reads) are indistinguishable, i.e., that the explicit JSD (or the normalized JSD, since they differ by a constant) between the models inferred from subsamples of these two samples was not higher than between data replicates of A, we used the following procedure. Without loss of generality, we assume that N ≤ K.

First, we split sample A into two non-overlapping samples A’ and A’‘, both of size N/2 sequencing reads. Then, we sample N/2 sequencing reads from B to obtain samples B’. After that, for a fixed subsample size S (S in [1000, 3000, 10000, 30000]) sequencing reads, we performed the following:

- Subsampling (without replacement) of S sequencing reads from the samples A’ and A’’ each thirty times independently – this yields thirty non-overlapping pairs and hence thirty (identically distributed) measurements of the JSD between data replicates of A (the explicit JSD of the models inferred from the samples within one pair).
- Subsampling of S sequencing reads from the samples A’ and B’ each thirty times independently – this yields thirty non-overlapping pairs and hence thirty independent measurements of the explicit JSD between subsamples of A and B (the explicit JSD of the models inferred from the samples within one pair).

We employed the null hypothesis that samples A and B are homogeneous, i.e., that the expected values of the RGMP inferred from them are equal. Consequently, the expected values of the RGMP inferred from subsamples of A’, A’’ and B’ are equal. Under the null hypothesis, the distance between model parameters values inferred from subsamples of A’ and A’’ will not significantly differ from the distance between model parameters values inferred from subsamples of A’ and B’. Hence, the mean of the distribution of the explicit JSD between A’ and A’’ models will not differ from that of A’ and B’ models. We used the unpaired Student’s T-test to check if the null hypothesis holds.

To investigate the properties of our test, we repeated the test procedure 30 times (Fig. S2A) for pre-B cell data for data replicates and for murine twin subjects (samples 1 and 2 from the MOUSE_PRE dataset) for the subsample size 3000 sequencing reads. We used the sample size 3000 due to the high computational cost of the procedure. In the first case (i.e. when the null hypothesis was known to hold), the p-value distribution resembled uniform distribution (Fig. S2B); in the second case, when the null hypothesis supposedly did not hold, the p-values were highly skewed towards zero (Fig. S2C).

## Software

IGoR v1.4.0 (Marcou, Mora, and Walczak 2018, <u>github</u>) for RGMP inference, nucleotide sequence Pgen evaluation and generation of synthetic AIRR-seq data. OLGA v1.2.3 (Sethna et al. 2019) for amino acid sequence Pgen evaluation. MiXCR v3.0.12 (Bolotin et al. 2015) for AIRR-seq data preprocessing and annotation.

## Graphics

Matplotlib v3.3.2 (Hunter 2007). Seaborn v0.11.0 (Michael Waskom et al. 2020).

## Hardware

Computations were performed on a dedicated server as well as the high-performance computing cluster FRAM (Norwegian e-infrastructure for Research and Education sigma2.no/fram).

## Data and code availability

The code used to generate the figures in this manuscript can be found in the following Github repository: https://github.com/csi-greifflab/desygnator, along with a link to the parameter files of all fitted IGoR models that were analyzed.

## Acknowledgments

The authors would like to thank Dr. Corey Watson, Dr. Gur Yaari, and Dr. Limenitakis for productive discussions and for their valuable feedback that has helped improve this study.

## Supplementary Material

**Supplementary Figure 1.**
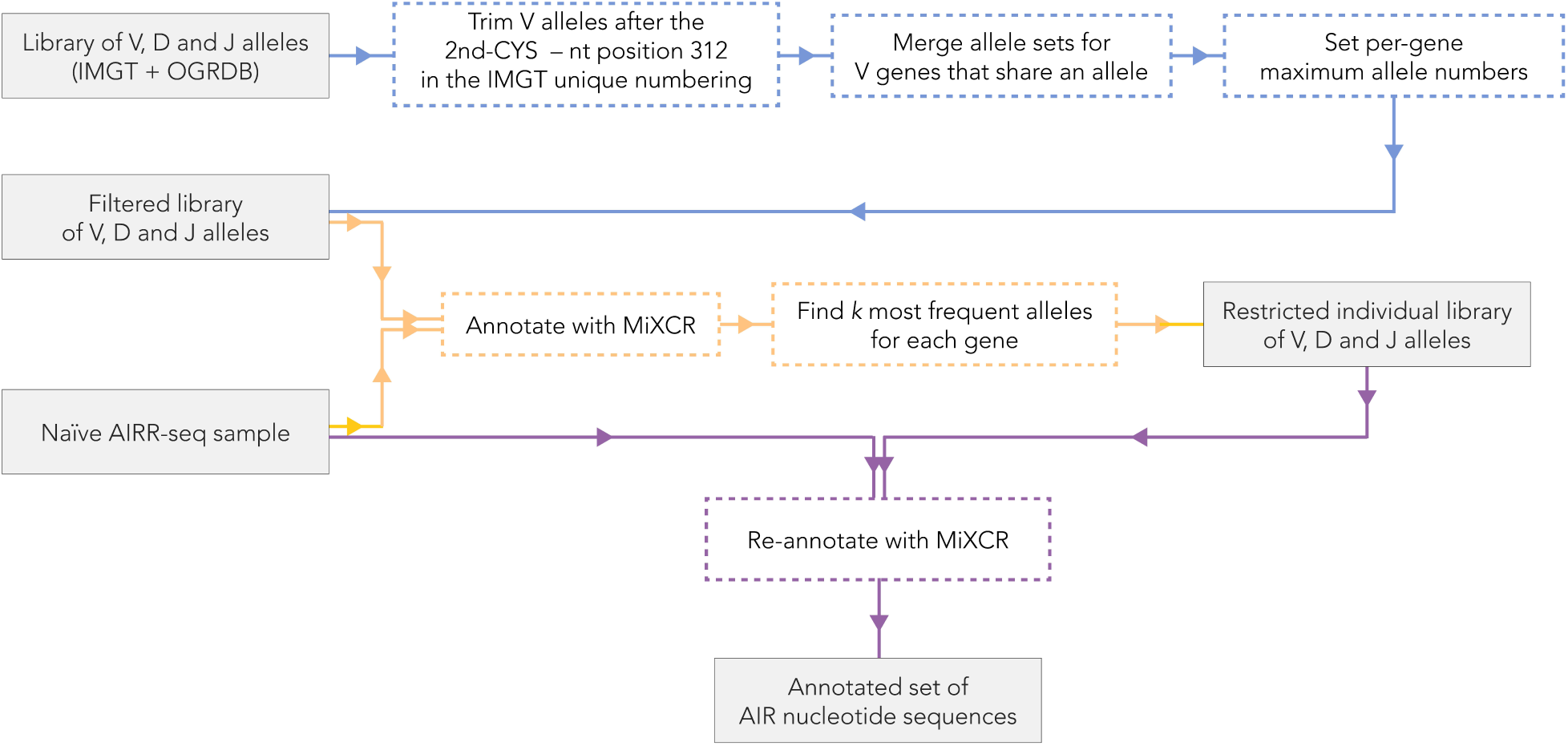
(relates to Fig. 6). Pipeline for allele-level annotation of AIR sequencing reads using validated allele sets deposited in publicly available databases. Blue path: preprocessing of the input library of IGHV alleles. Orange path: computation of an individually restricted allele library for a given AIRR-seq sample. Violet path: annotation of AIRR sequencing reads in the sample using the individually restricted allele library.

**Supplementary Figure 2.**
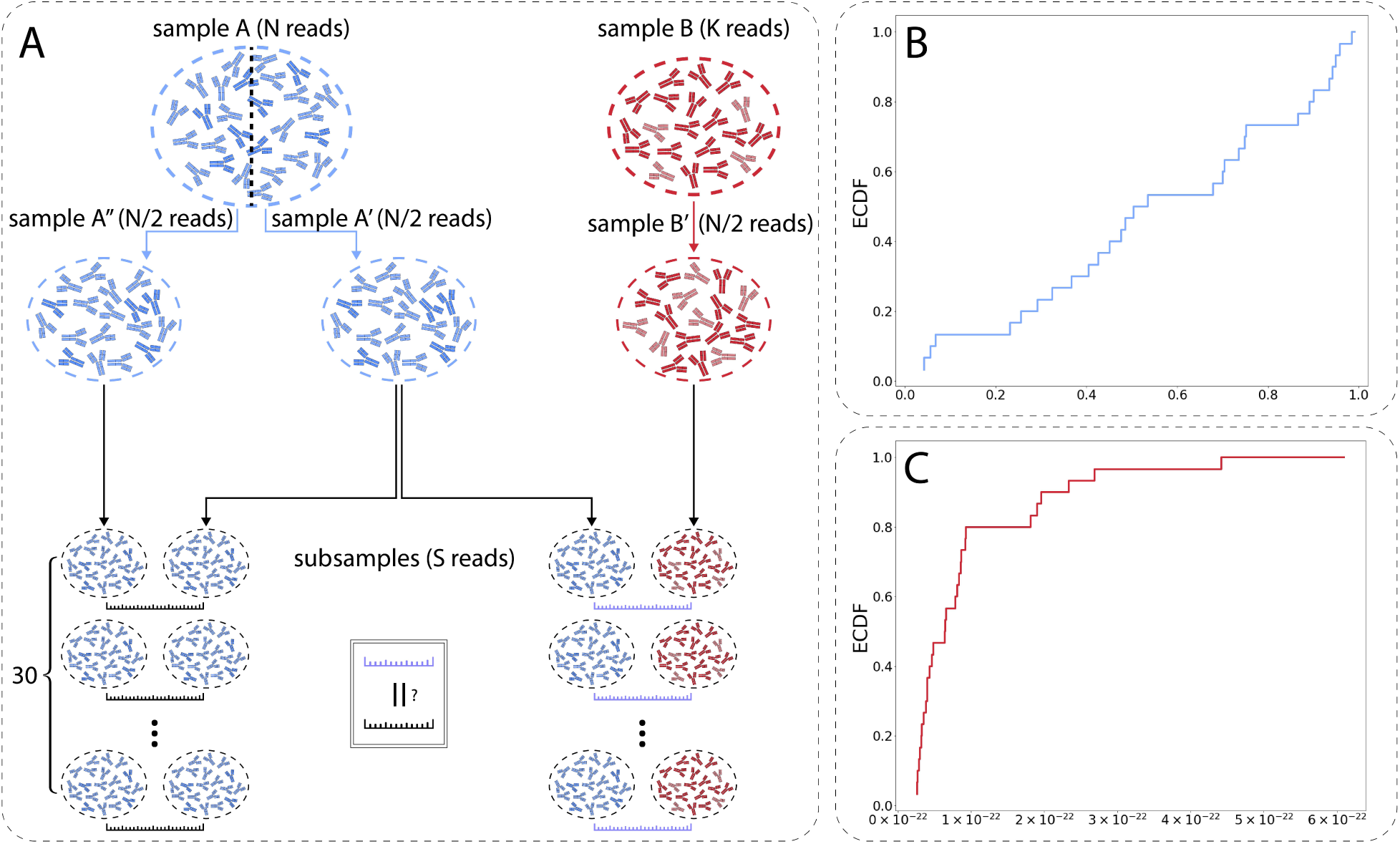
(A) Subsampling from a pair of experimental samples to obtain the 30 independent measures of the explicit JSD values for the sample size of S sequencing reads (S in [1000, 3000, 10000, 30000]). (B) In the case when the null hypothesis holds (A and B are data replicates), for the sample size of 3000 sequencing reads, p-values of the Student’s T-test for repertoire comparison replicated 30 times are distributed uniformly between 0 and 1, as shown by the empirical cumulative distribution function. (C) In the case when the null hypothesis supposedly does not hold (A and B are samples obtained from twin subjects), for the sample size of 3000 sequencing reads, the p-value distribution is highly skewed towards 0.

**Supplementary Figure 3.**
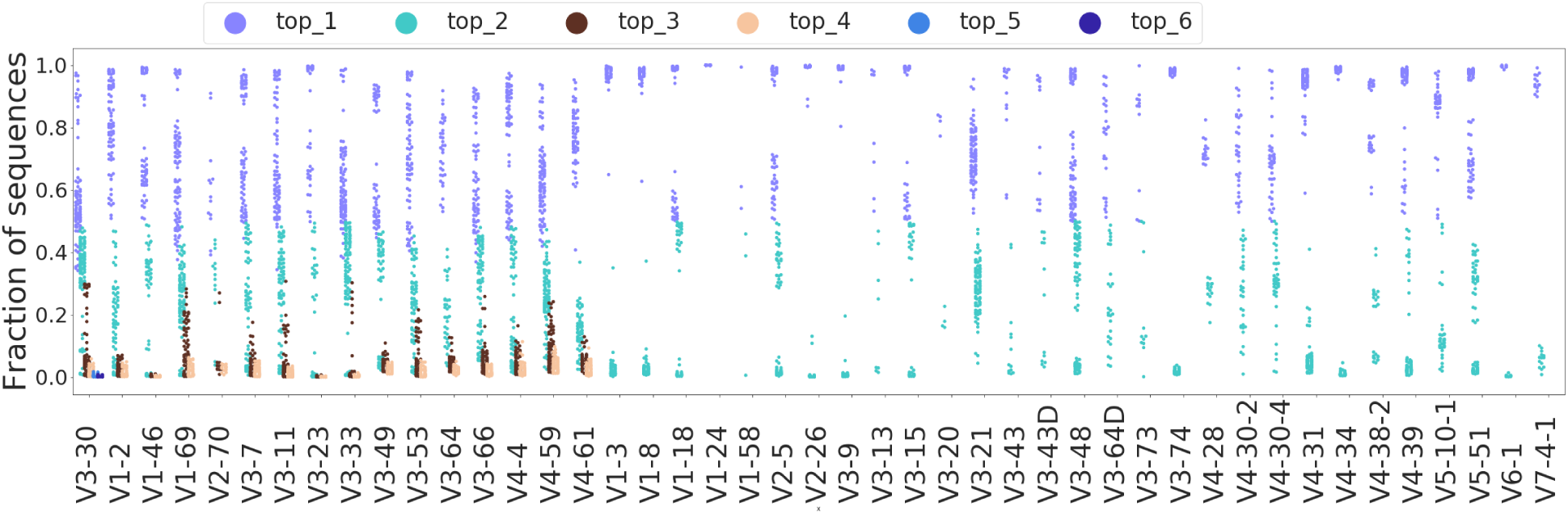
(relates to Fig. 6). Allele fractions in the HUMAN3 dataset after applying the pipeline described in the Methods 3, similar to Fig. 6G. The yellow points corresponding to the fractions of the remaining (i.e., not among the top_k) alleles were redistributed across other colors – as each of the misaligned sequencing reads was re-aligned to one of the top_k alleles.

**Supplementary Figure 4.**
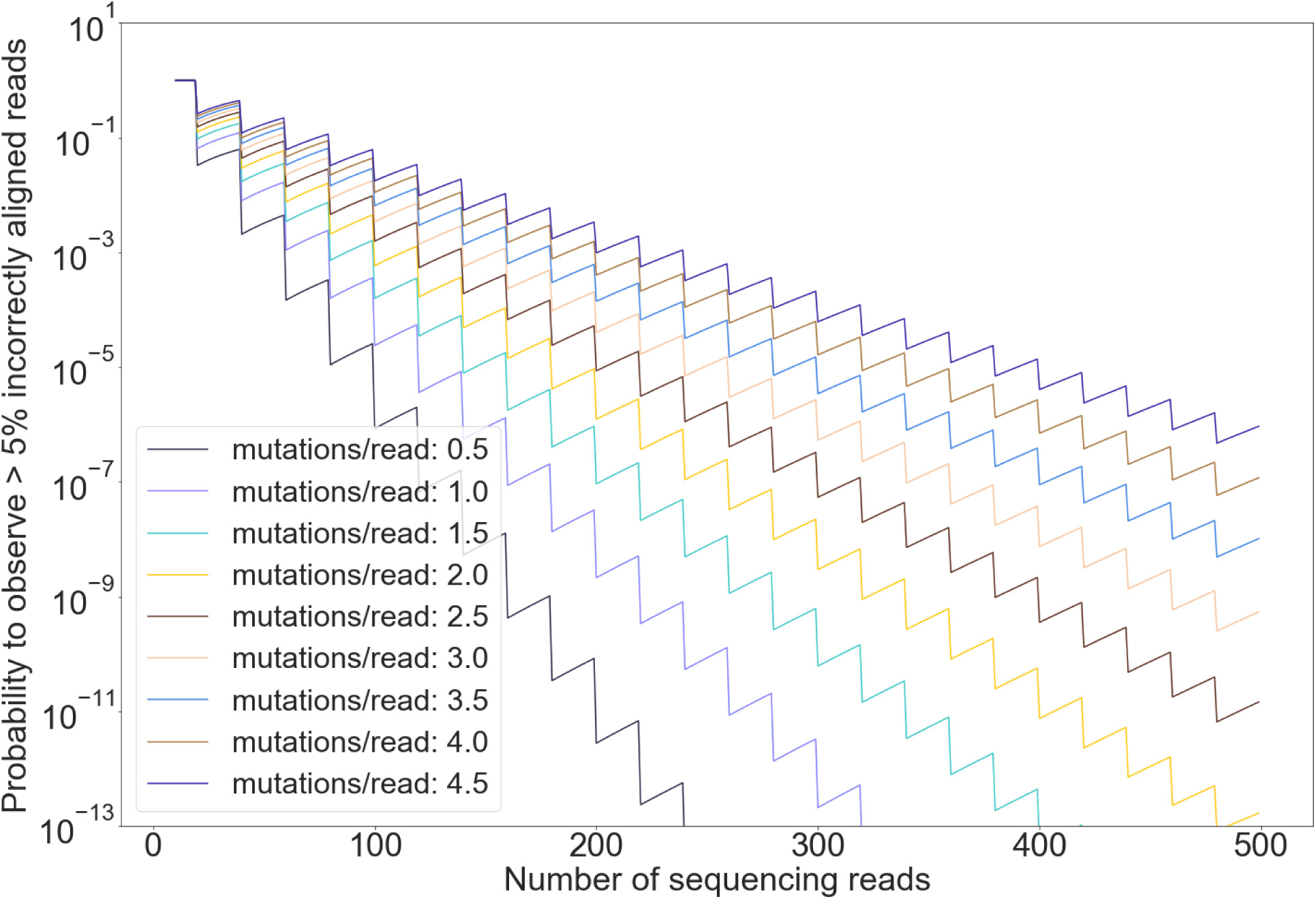
(relates to Fig. 6). The theoretical probability to observe more than 5% incorrect allele assignments for reads originating from a given V segment. The probability is calculated for the worst-case scenario under the following assumptions: (i) there are two alleles that differ by a single nucleotide substitution on position *Pos_x.* (ii) nucleotide substitutions in the sequencing reads are independent for different positions (as we talk about naïve B cell data and we can be sure that these substitutions are caused by sequencing/PCR errors and not somatic hypermutations), (ii) the number of errors per read is below 4.5 (Wardemann and Busse 2019). In such conditions, the probability to misalign a read is the probability for the nucleotide on position *Pos_x* to be mutated. Since two alleles differing by a single nucleotide substitution is the worst case, the calculated probability is an upper bound for a probability to misalign an arbitrary sequencing read. If there are sequencing reads in the sample and the probability to observe a SHM on a given position is *p_mut_* (which is proportional to the average number of mutations per sequencing read), then the probability to observe more than 5% of misassigned alleles is 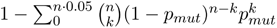. The zigzag shape of the plot is due to the number of terms in the sum in the formula: every time the number of reads increases by twenty, the upper limit for increases by one. The calculations show that the threshold of 500 reads ensures a reliable probability of observing less than 5% misalignments even for the upper bound of the error rate.

**Supplementary Figure 5.**
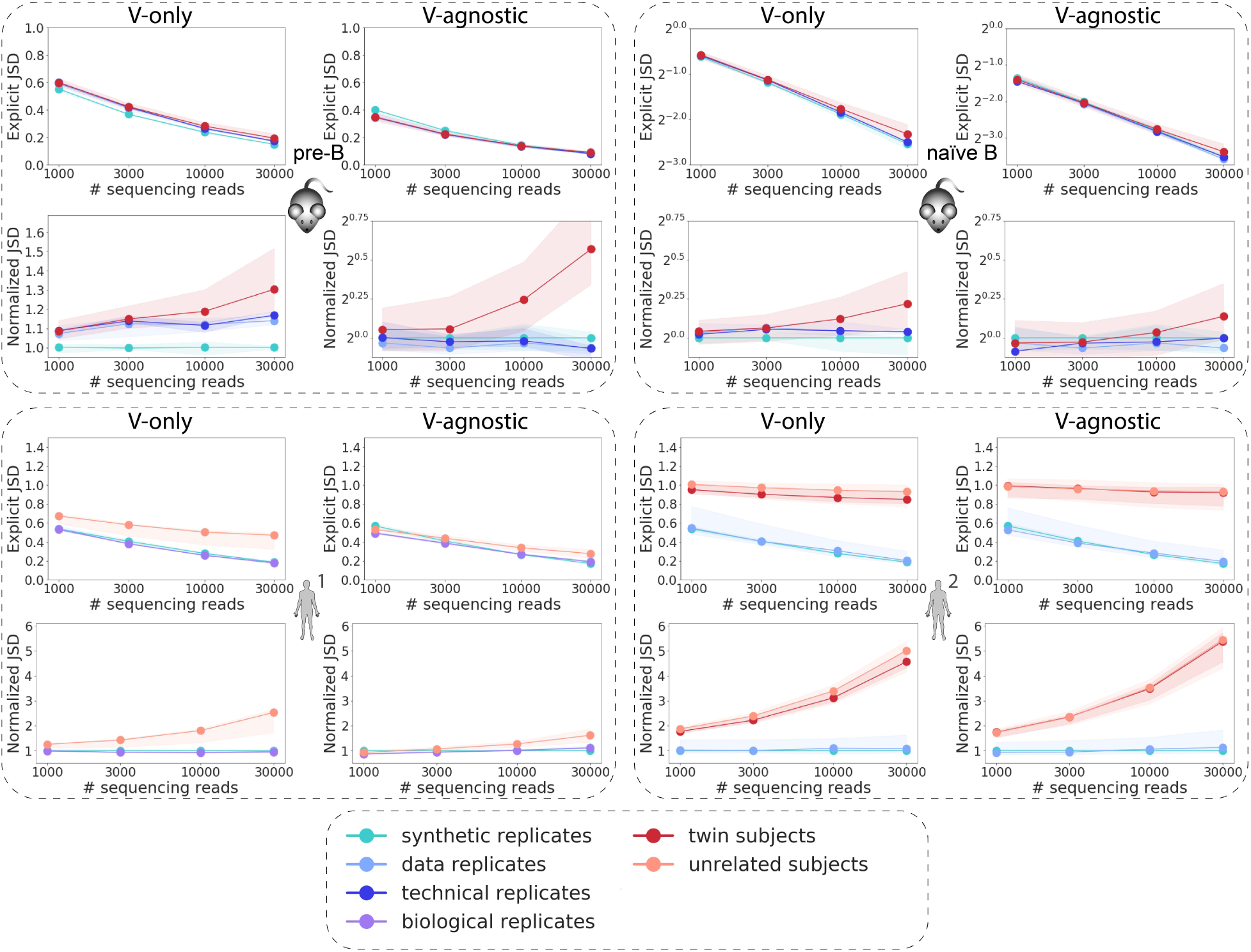
(relates to Fig. 2). JSD-based distance for V segment-only and V segment-agnostic explicit and normalized JSD. Analogous to Fig. 2 except for here the JSD was computed either using only the V choice probability distribution and V deletion profiles (columns 1 and 3) or in a V segment-agnostic way (columns 2 and 4) for MOUSE_PRE (upper left), MOUSE_NAIVE (upper right), HUMAN1 (lower left) and HUMAN2 (lower right) datasets.

**Supplementary Figure 6.**
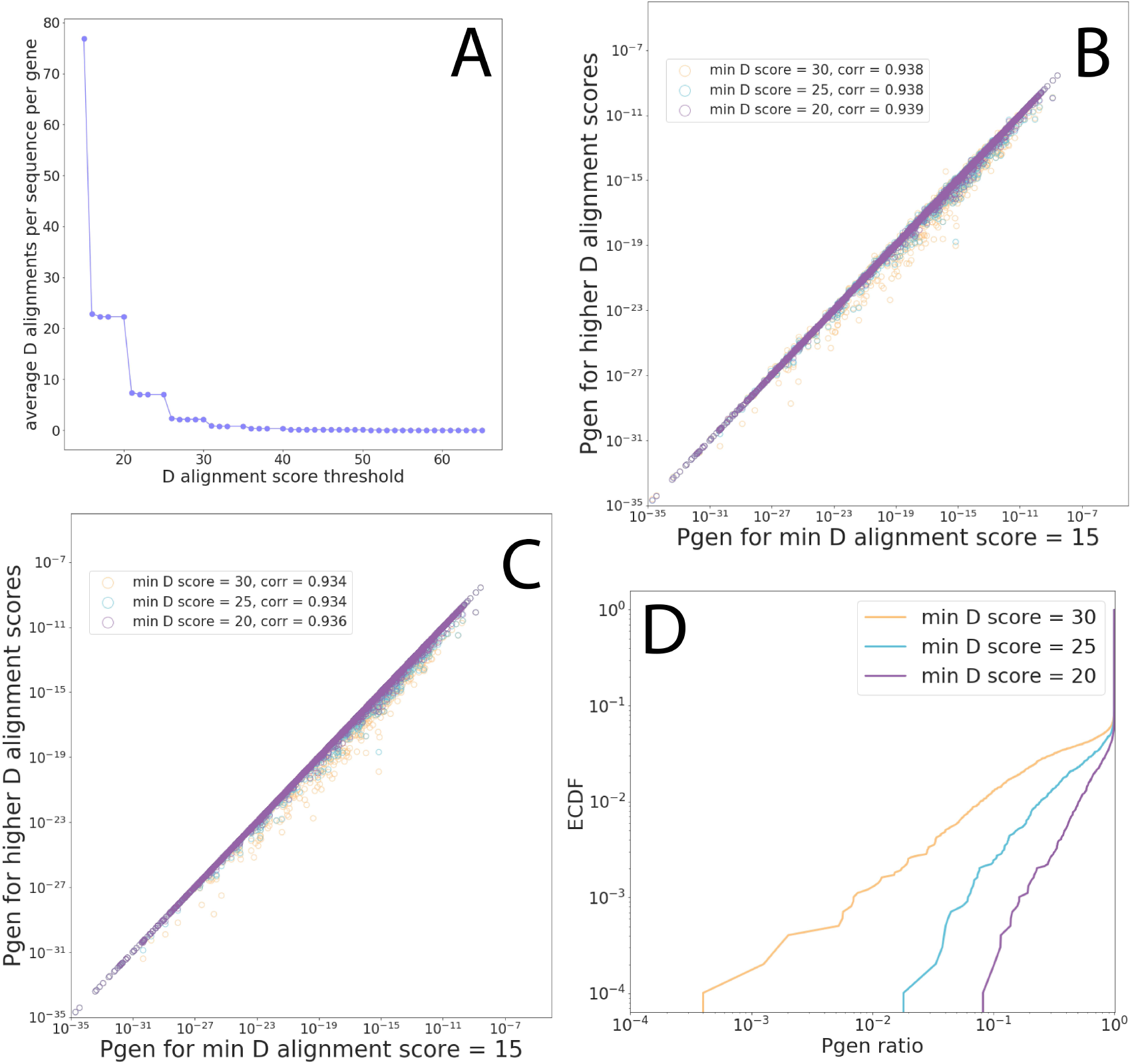
(relates to Fig. 6). IGoR behavior with different D gene alignment score thresholds. Here, we show the argumentation for increasing the D gene alignment score thresholds in IGoR from 15 to 30. (A) A synthetic dataset of 10000 sequencing reads generated using ImmuneSim was aligned using IGoR. With the default value for D gene alignment score threshold (15), IGoR produces an enormous number of alignments: more than 80 alignment variants per sequence per gene on average. We filtered the alignments, leaving only alignments with scores higher than 20 (violet), 25 (blue) and 30 (yellow). Filtering the alignments reduces the number of alignment variants per sequence per gene to the more realistic 8 (for threshold=25) or 2 (for threshold=30). (B) After that, we inferred the RGMP for all four variants of alignments and used the four inferred RGMP sets to evaluate the sequence Pgens. To plot the Pgens estimated using the modified alignment score threshold as a function of the ones estimated using the default threshold, we showed them on a scatter plot: X-axis stands for the D gene alignment score threshold = 15, Y-axis stands for the higher ones. The Pgens are highly consistent (Pearson’s r > 0.93), suggesting that one can use IGoR with an increased D gene alignment score threshold. Out of 10000 sequencing reads, 107 Pgens equaled zero (108 for threshold=30). (C) The same as B, but all Pgens were computed using the same model (inferred from alignments computed with the default parameters) but using different alignments during the evaluation step. The Pgens are also consistent in this case (Pearson’s r > 0.93). (D) The Pgen ratio distribution in a log-log scale (Pgens for higher score thresholds divided by Pgens for threshold=15).

**Supplementary Figure 7.**
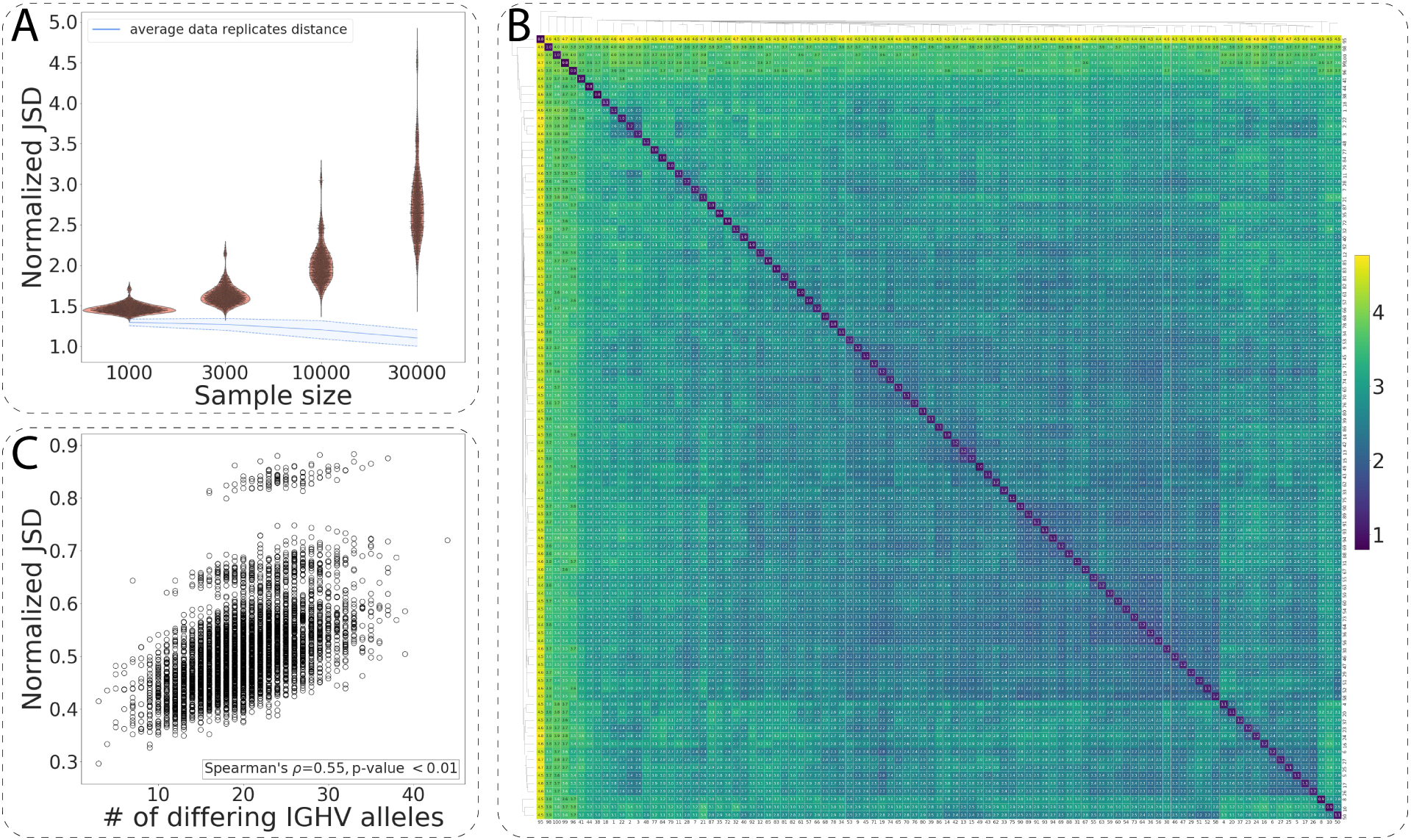
(relates to Fig. 3). The normalized V segment-only JSD applied to the HUMAN3 dataset. Analogous to Fig. 3 but here the normalized JSD was calculated using only the V segment-related components (V segment choice and V deletion). The results support those obtained with the full model (Fig. 3).

**Supplementary Figure 8.**
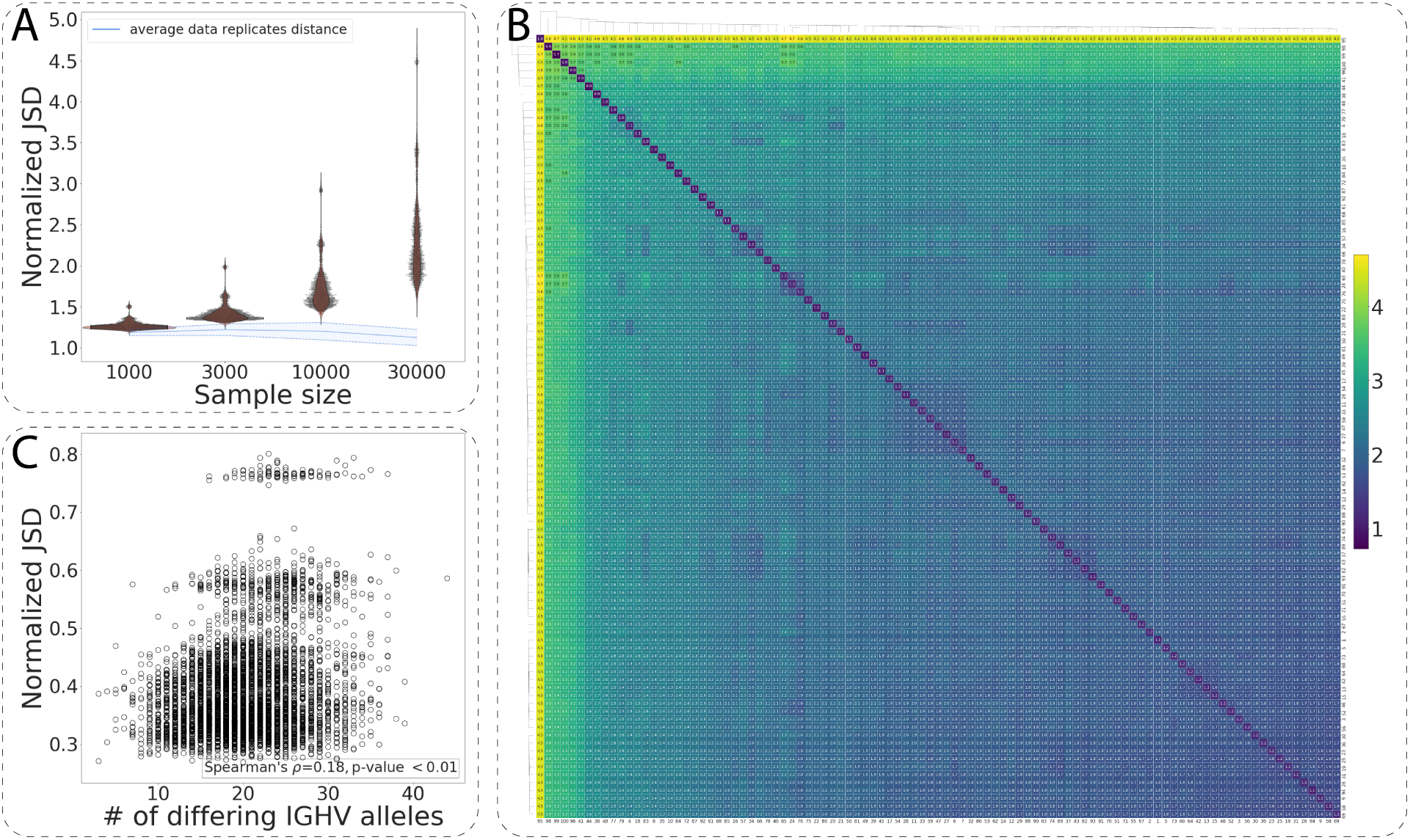
(relates to Fig. 3). The normalized V segment-agnostic JSD applied to the HUMAN3 dataset. Analogous to in Fig. 3 but the normalized JSD was calculated only using V segment-agnostic components. The results support the ones obtained with the full model (Fig. 3).

**Supplementary Figure 9.**
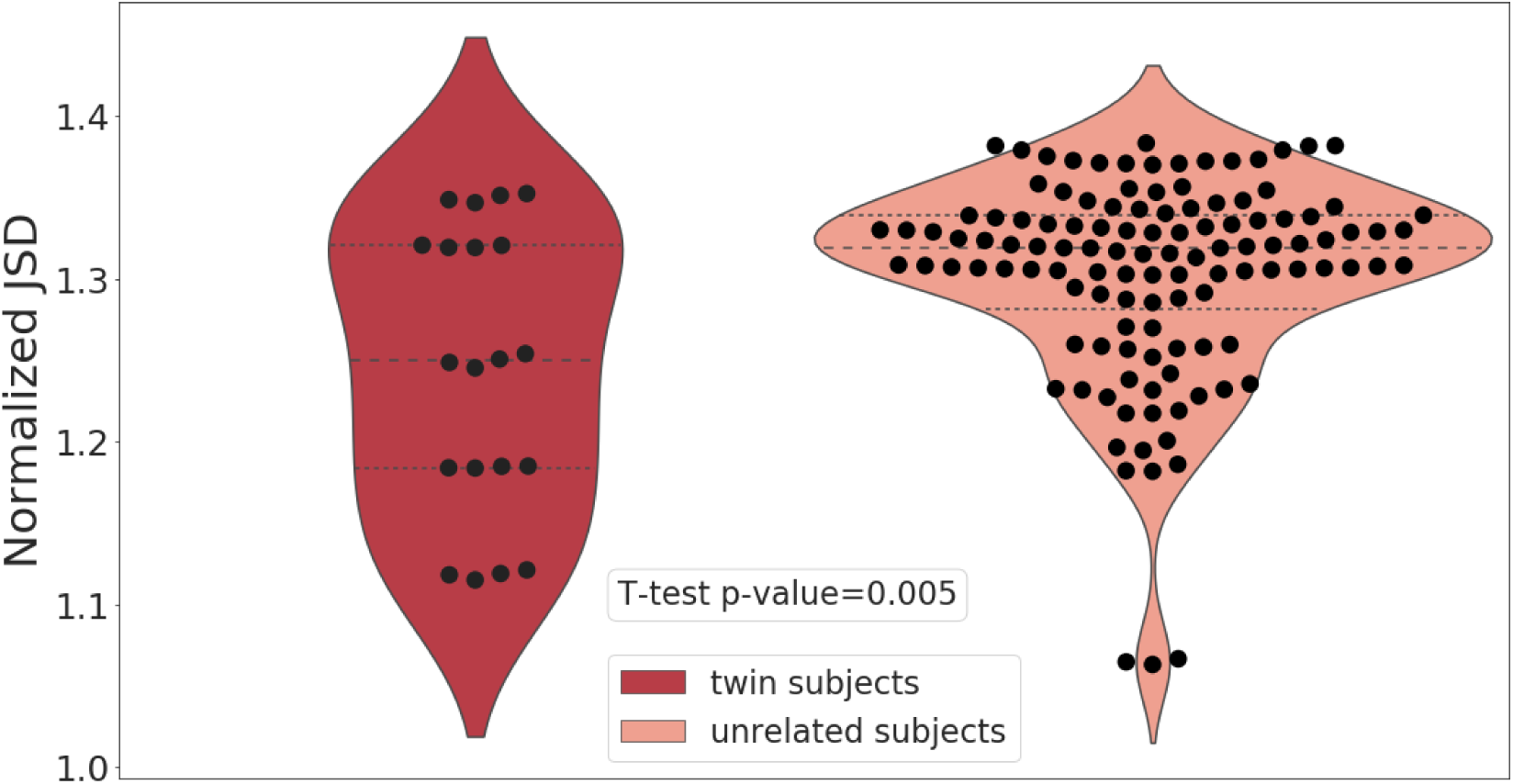
(relates to Fig. 2). The normalized JSD distance for five twin pairs from the HUMAN2 dataset. The normalized JSD within the pairs of twins (left) and across the pairs of twins, i.e., for unrelated individuals (right) in the HUMAN2 dataset, sample size=30000 sequencing reads. The dashed lines correspond to the quartiles of the normalized JSD distribution. The normalized JSD for unrelated individuals is on average higher than the normalized JSD between the twins. In some cases, the normalized JSD for twins is lower than for unrelated individuals: we speculate that this may be because genetic factors account for only a small portion of the JSD and, consequently, variation in non-genetic factors may be stronger than the impact of genetic factors.

**Supplementary Figure 10.**
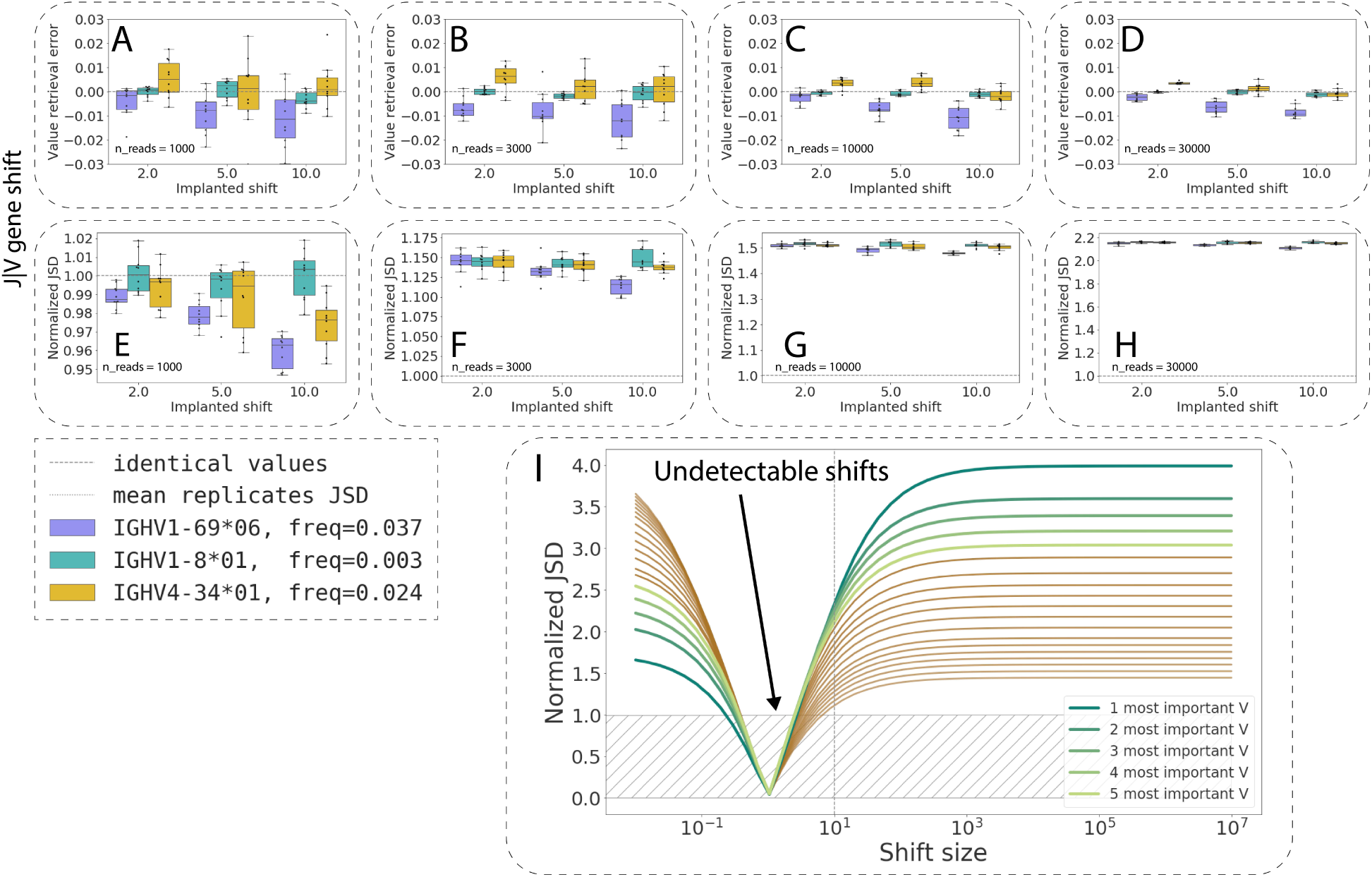
(relates to Fig. 5). Sensitivity of IGoR and the normalized JSD to the V choice probabilities. Identical to Fig. 5 but computed for the V choice parameter (instead of the J|V conditional choice parameter).

